# Transposable element landscape changes are buffered by RNA silencing in aging *Drosophila*

**DOI:** 10.1101/2021.01.08.425853

**Authors:** Nachen Yang, Satyam P. Srivastav, Reazur Rahman, Qicheng Ma, Gargi Dayama, Madoka Chinen, Elissa P. Lei, Michael Rosbash, Nelson C. Lau

## Abstract

Genetic mechanisms that repress transposable elements (TEs) in young animals decline during aging, as reflected by increased TE expression in aged animals. Does increased TE expression during aging lead to more genomic TE copies in older animals? To answer this question, we quantified TE Landscapes (TLs) via whole genome sequencing of young and aged *Drosophila* strains of wild-type and mutant backgrounds. We quantified TLs in whole flies and dissected brains and validated the feasibility of our approach in detecting new TE insertions in aging *Drosophila* genomes when natural defenses like RNA interference (RNAi) pathways are compromised. By also incorporating droplet digital PCR to validate genomic TE loads, we confirm TL changes can occur in a single lifespan of *Drosophila* when TEs are not suppressed. We also describe improved sequencing methods to quantify extra-chromosomal DNA circles (eccDNAs) in *Drosophila* as an additional source of TE copies that accumulate during aging. Lastly, to combat the natural progression of aging-associated TE expression, we show that knocking down *PAF1*, a conserved transcription elongation factor that antagonizes RNAi pathways, may bolster suppression of TEs during aging and extend lifespan. Our study suggests that RNAi mechanisms generally mitigate genomic TL expansion despite the increase in TE transcripts during aging.

## INTRODUCTION

All animal genomes carry the genetic burden of a sizeable reservoir of parasitic elements called transposons or transposable elements (TEs). This TE burden can range from the extreme >70% proportion of the axolotl genome [1, 2] to >50% in the human genome [3] to >10% in the *Drosophila melanogaster* genome [4, 5]. TEs are selfish invaders of animal genomes with some potential for stimulating more rapid gene regulatory innovations like serving as novel enhancers [6], but more frequently are detrimental to animal fitness when they insert into and disrupt expression of important genes [7]. Therefore, conserved chromatin regulation and RNA-interference (RNAi) pathways must silence TEs to ensure fertility and animal health. However, these genomic defense mechanisms also weaken during animal aging concomitant with observable decreases in genomic integrity in aging cells. This phenomenon has been articulated in the hypothesis of TEs impacting aging [8].

Initial support for this hypothesis in the model organism *D. melanogaster* came from studies of TE expression increasing in aging flies [9–11, 12]. For example, mutants in chromatin silencing factors and RNAi pathway genes which repress TEs have reduced lifespans [9, 10, 13–16], whereas dietary restriction and overexpressing the RNAi and chromatin factors can limit TE expression and promote longevity [13, 14]. Neurodegeneration modeled in aging flies through overexpressing aggregating proteins like TDP-43 and TAU also leads to elevated TE expression [17–20]. Additionally, there is evidence of a somatic population of Piwi proteins which can serve an additional TE defense mechanism that when mutated leads to shorter lifespan and loss of stem cell maintenance [13, 16, 21–23].

Beyond flies, mammals also must repress TEs for critical development of germ cells, embryos and neurons. Mammals have a complex, interconnected network of silencing pathways like the axis of SETDB1 [24, 25], KAP1 [26–28] and the HUSH complex [29–32]; and its cooperation with histone deacetylases like SIRT6 [33, 34] and histone methyltransferases like Suv39h1 and G9A [35–37]. In addition, there are DNA methyltransferases that genetically interact with the piRNA pathway to target TEs for chromatin silencing in mammalian germ cells [38–44]. In primates and mice, the most active TE is the *LINE-L1* which is implicated in somatic genome mosaicism in developing brains and individual neurons [45–53]. Lastly, *LINE-L1* is linked to deleterious novel mutations in tumors and they are activated in cell culture models of cellular senescence [54–59]. Although TE control is clearly important to mammalian health, the large genome sizes and longer lifespans hampers comprehensive assessments of mammalian TLs during impact aging.

Therefore, in this study we leveraged *Drosophila’s* rapid aging, its compact genome and powerful genetic tools as significant advantages for testing how TE landscapes may change during normal animal aging. An important goal of our study is to address the debate of whether TLs quantitated from Whole Genome Sequencing (WGS) of *Drosophila* genomes represent true gains in TE genomic load [60]. One bioinformatics program called TEMP [61] has been used extensively in determining TE insertions from *Drosophila* WGS [62, 63] but its capacity to distinguish bona fide TE insertions from potential library sequencing artifacts has been re-examined [60]. Noting the high degree of variability in TE insertion calls from various bioinformatics programs applied to *Drosophila* WGS data [64], we therefore developed our own program called the Transposon Insertion & Depletion AnaLyzer (TIDAL) to identify the tremendous diversity of TLs across various *Drosophila* strains [65]. TIDAL’s increased specificity in TE determinations comes from requiring sequencing reads mapping to both sides of genomic locus flanking the TE insertion. This specificity was benchmarked against genomic PCR tests [65], and TIDAL has characterized TLs in other *Drosophila* studies of genetic factors regulating TE silencing [66, 67].

In this study, we demonstrate how WGS and extrachromosomal circular DNA (eccDNA) sequencing of aged and young flies can report changes in TLs during fly aging. Although TE RNA upregulation is a recurring phenotype of aging wild-type flies, we show that major changes in the genomic TLs are generally suppressed by the RNAi pathway because RNAi mutants allow TEs to expand their genomic DNA (gDNA) copy numbers. We also demonstrate that tissue-specific (i.e. fly brain tissues) gDNA sequencing can sensitize the detection of genomic TL changes; and eccDNA accumulation during aging of the *ISO1* strain is an additional feature of the hypothesis of TEs impacting animal aging. Lastly, we show that genetically boosting RNAi activity in aged flies via knockdown of *PAF1* can suppress TE RNAs and extend longevity. Together, these results demonstrate that the RNAi pathway buffers genomic alterations by the natural increase of TE RNAs during aging and suggest *PAF1* inhibition in aging animals could be a therapeutic target in this genetic mechanism of TE repression.

## RESULTS

### Recurring increase of TE RNA expression during fly aging

Although previous studies using certain control wild-type (WT) fly strains showed that TE RNAs were upregulated in aged flies [10, 14], we decided to reconfirm this observation for three commonly used WT fly strains that would form the basis of this study. Using our lab’s standard rearing conditions, we first determined the aging curves for the *ISO1* strain used for the *D. melanogaster* reference genome sequence [4], an isogenic *w1118* strain that is a common background strain in genetic studies [68], and the *Oregon-R* strain used in a series of functional genomics datasets [69]. Whereas *w1118* and *OreR* displayed lifespans typical of other WT fly strains (**Figure 1A**), the shorter lifespan of *ISO1* was expected because its genetic background was known to sensitize phenotypes from chemical mutagenesis screens [70].

**Figure 1.**
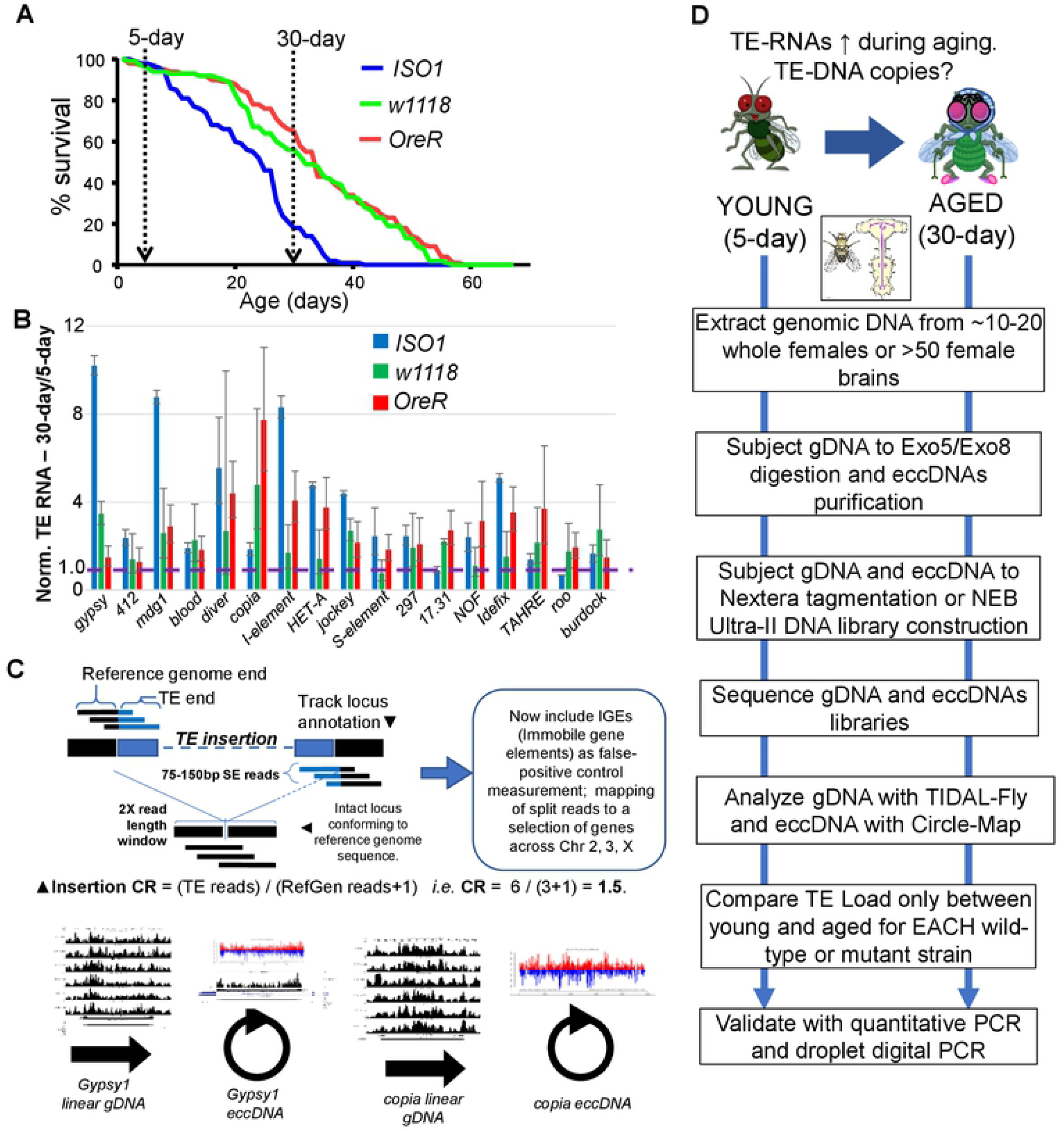
Overview of study to examine whether TE-DNA copy numbers change during fly aging. (A) Survival curves of the three wild-type fly strains carried out in this study, indicating the selection of 30-day adults as a representative timepoint of aging onset. (B) Validation of TE transcript expression increases during fly aging through qRT-PCR of TE RNAs normalized to *rp49* transcripts. Error bars are propagated standard deviations of delta-CT values from three replicates. (C) Overview of TE detection strategy from Whole Genome Sequencing (WGS) data using updated TIDAL-Fly and extra-chromosomal circular DNA (eccDNAs) detection scripts. (D) Study designed for comparing TE load between 5-day young and 30-day aged flies within each wild-type and mutant strain.

We then followed the experimental convention of other studies [10, 14] to standardize the comparison of 30-day aged adults versus 5-day young adults, and we performed quantitative RT-PCR on a panel of TEs from total RNAs from females (Fig. 1B). We replicated many examples of TE RNAs being upregulated in the aged WT flies but noticed variability in which specific TE families were the most significantly upregulated during aging. For example, *gypsy, mdg1*, and *I-element* were up regulated at the RNA level in *ISO1* aged flies, while *copia* and *1731* RNAs were upregulated in *w1118* and *OreR* (Fig. 1B). This variability may reflect the inherently distinct TLs between these three strains [65], but the trend holds true that WT adult flies recurringly experience increased TE expression during aging.

However, some previous studies examining *Drosophila* TEs during aging mainly used a genetic reporter called the *gypsy-TRAP* to reflect increased transposition activity in aging flies [13, 14, 18, 71]. This reporter has the advantage of low cost and sensitivity of detecting small numbers of cells in a background of nonmodified cells, yet this transgenic construct is also only designed for *gypsy* to insert and activate a fluorescent protein read-out and cannot assess overall TLs. A newer and distinct TE activity reporter called the *gypsy*-CLEVR puts the fluorescent protein expression cassette into the domain of a 5X-UAS promoter only after retrotransposition, to take advantage of cell-type or tissue specific GAL4 driver *Drosophila* strains [15]. Only one recent study we are aware of assessed TLs during fly aging by WGS of enriched αβ-Kenyon Cell neurons [60] and which argued that various pitfalls obscured the ability to observe TL increases during fly aging. For example, the study itself discussed that Multiple Displacement Amplification (MDA) required to amplify the minute amount of neuronal gDNA prior to Illumina library construction could contribute to artifactual chimeric molecules that represent false positive TE insertions [60]. Therefore, our more comprehensive effort to examine TLs through direct WGS should add valuable insight to this question.

First, some issues need to be considered in TL determinations in *Drosophila* WGS datasets, such as two different TE-insertion discovery programs, TEMP [61] and TIDAL [65] that each can yield different results from analyses of the same dataset (**Figure S1**). Balancing sensitivity against specificity, TIDAL has similar trends as TEMP in revealing the diversity of TLs amongst *Drosophila* samples (Fig. S1B) and both are effective at calling germline insertions, but TIDAL avoids false positive predictions that others have contended as somatic TE insertions (Fig. S1C, [60]). TIDAL handles this issue differently by computing a Coverage Ratio (CR) score for each TE insertion from pooled sequencing of a small group of flies (Fig. 1C), where TE insertion reads are divided by reference genome mapping reads and a pseudocount of 1; such as a CR of 2 that we used as an arbitrary cutoff for indicating deep penetrance of a TE insertion at a given insertion locus. When we re-analyzed the WGS datasets from [60] with TIDAL even with the caveats of MDA, the TIDAL outputs do suggest increasing TLs within the fly aging neuron genomes (**Figure S2**).

### Measuring TL differences by direct WGS and the TIDAL program

To meaningfully compare TL changes during a single generation of aging flies from WGS and to avoid the genomic complications of normalizing against Y-chromosome reads that are exceptionally dense with repeats [72], we only compared samples from within the same strain in small numbers of young versus aged whole female adult flies or female brains (Fig. 1D). In our process we extracted a set amount of genomic DNA from 10 flies that allowed for reproducible WGS library construction without requiring MDA or other total DNA amplification methods. We then sequenced on the Illumina platform each fly strains’ bulk gDNA library to a minimum >~30 million 75-bp reads for >~16X fold genomic coverage of the version Dm6 genome assembly (**Table S1**). Each library was analyzed identically with the TIDAL program [65] and new TE insertions were counted individually and then normalized against the reads per million measurement to account for sequencing depth differences. In developing our own methodology to examine fly TLs during aging, we recall our previous study showing that each fly strain’s unique TLs depends on how inherently distinct its genetic background is from the reference genome strain *ISO1* [65]. Therefore, it was expected that new TE insertions quantified and normalized against each library’s sequencing depth would yield the lowest numbers for *ISO1* and the most TE insertion differences in *w1118* and *OreR* (**Figure 2A**).

**Figure 2.**
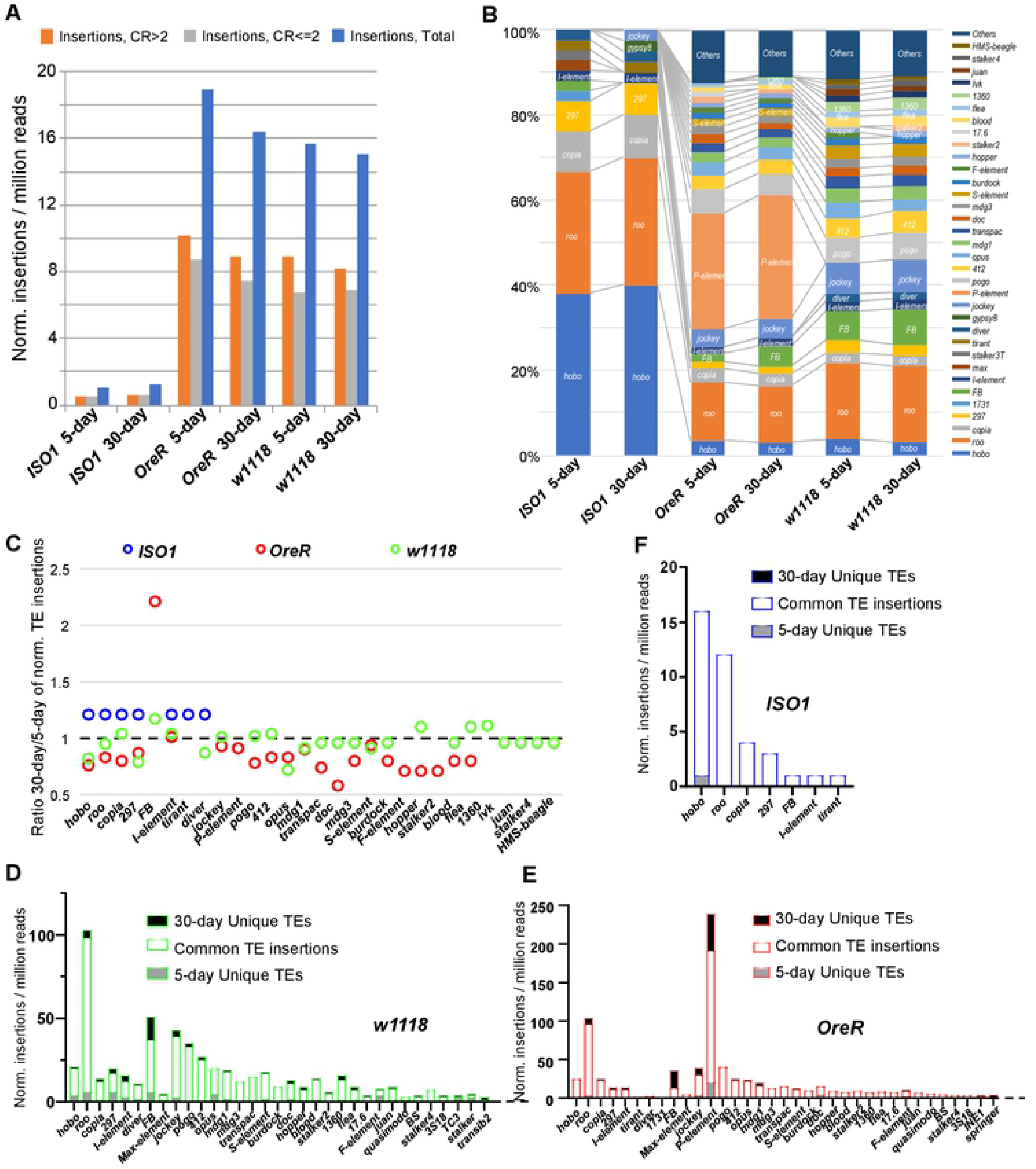
WGS analysis of TE insertion numbers between 5-day young versus 30-day aged wild-type fly strains. (A) Quantification of new TE insertions as compared to the reference genome using the TIDAL-fly program. Categories of total TE insertions broken by the Coverage Ratios (CR) of CR>2 and CR<=2. (B) Within each strain, TE families’ percentages are ordered by the color legend. (C) Ratios of the 30-day versus 5-day of normalized TE insertions from panels D-F. Note the *w1118* strain has the greatest number of distinct TE insertions detected by TIDAL. (D-F) Number of unique TE insertions (filled bars) present in 5-day and 30-day relative to common insertions present in both samples (open bar) of *w1118, OreR* and *ISO1* fly strains. These panels display only the TE families that were detected by TIDAL be at least 1% of total number of TE families (*i.e*. all the TEs not lumped into the “Others” category of Fig. 2B).

As expected, each of these WT flies TLs displayed completely distinct compositions of new insertions of TE families relative to the Dm6 reference genome sequence (Fig. 2B), such as a larger proportion of *hobo* TEs in *ISO1*, major infiltration of *P-elements* in *OreR*, and several more *FB, pogo* and *412* TEs in *w1118*. Focusing on the ratio of 30-day to 5-day insertions for the specific TE families making up the bulk of these strains TLs, we could observe some increases in TE insertions in aged flies as well as decreases (Fig. 2C). This was reflected at the total TL level with modest new TE insertions in 30-day aged *ISO1* flies versus 5-day young flies, with also perplexing total decreases in *OreR* and *w1118* flies (Fig. 2A). Although the vast majority of the TE insertions were commonly detected by TIDAL in both 5-day and 30-day *w1118* and *OreR* flies (Fig. 2D,E), there were more TE insertions only detected in these 5-day young fly genomes than of 30-day aged flies. Only a few *hobo* insertions were also only seen in 5-day young *ISO1* flies and were no longer detected in 30-day aged flies (Fig. 2). This observation can be explained by this analysis that only focuses on TE insertion counts as quantile samplings of reads discordant from the reference genome. Thus, a new somatic transposition event in a small subset of cells could be overshadowed by a background of unmodified reference sequences and could explain a TE with a low CR score that is sampled in 5-day fly gDNA sample but then missed in the 30-day sample. This is a known limitation of the WGS approach and sacrificing sensitivity to improve specificity in the original TIDAL program [65].

Therefore, we updated TIDAL to map to *Drosophila* TE families consensus sequence coverage, and added arbitrarily-selected protein coding genes, analogous to the modification to TEMP to track protein-coding genes as Immobile gene elements (IGEs) [60]. We gauged a relatively low average rate (<4%, Table S1) of false positive split reads called by TIDAL in hitting IGEs, whereas these protein coding genes sequencing coverage generally also remained stable between 5-day young and 30-day aged flies. (**Figure S3A**). Tracking TE consensus sequence coverage has the advantage of accounting for all accumulating TE sequences in both the mappable and unassembled and ambiguous-mapping regions of the genome. With this analysis approach, we could detect a clearer increase of total TE sequence coverage in *OreR* and *w1118* 30-day aged flies versus 5-day young flies (Fig. S3A). This overall aging-associated increase in TE consensus sequence coverage was represented by many TE families and is controlled against stable protein-coding gene coverage (Fig. S3B). However, both this coverage analysis and the quantile insertions analysis cannot discriminate between a full-length or truncated TE sequence, which we have noted in *P-elements* can have critically variable transposition activities [73].

### Resolving and validating our approaches measuring TE landscapes with RNAi mutants

This unresolved genomics challenge of using short read WGS data for analyzing TE sequences coverage also extends to some limitations in using droplet digital PCR (ddPCR) to precisely quantify genomic TE copies for only the isoforms covered by the short ddPCR amplicons [45]. Although the ddPCR quantifications of specific TE copies (Fig. S3C) followed the similar proportional trends of TE families called by TIDAL (Fig. 2B), the aging-associated increases in TE copies measured by ddPCR in the WT fly strains was also not detected. In questioning the accuracy of this ddPCR assay in absolute quantification of TE copies, we compared ddPCR results on *P-elements* and *I-elements* versus WGS and TIDAL determinations in two other directly matched gDNA samples (**Figure S4A,B**). The ddPCR copy number measurements were very similar to the WGS and TIDAL determinations, indicating both methodologies are consistent with each other in the quantifications. Furthermore, we replicated a previously reported genetic cross [74] that in one format triggers a large burst of *I-element* transposition in the embryos but in a second format maintains *I-element* silencing (Fig. S4C). We reanalyzed the WGS datasets from [74] with TIDAL reporting 505 new *I-element* copies versus the 3732 insertions called by TEMP in that study, with our ddPCR results leaning closer to the TIDAL count (1590 copies, Fig. S4D,E). These data reaffirm the findings from [74] that the oocyte is the critical battleground between the host and the selfish genetic element.

To explain why aging-associated TL changes seemed muted or were challenging to detect in WT fly strains, we considered two competing hypotheses: (1) non-penetrant TE insertions are masked by multiple unmodified genomic loci within the pools of sequences imposing limitations in WGS and TIDAL analysis versus (2) WT flies retain RNAi defenses like TE-targeting siRNAs [21, 22, 75–77] and piRNAs [13, 78–80] to prevent increasing TE RNAs from completing genomic transposition events. To test these hypotheses, we collected the same 5-day and 30-day aging cohorts from three sets of different mutants in the two main arms of the RNAi pathway in *Drosophila* (**Figure 3**). We analyzed two independent mutants each in the *piwi, aubergine, and AGO2* genes and conducted the same whole flies WGS and TIDAL analysis as the WT strains. In each of these six mutants, TLs showed dramatic increases in new TE insertions during fly aging (Fig. 3A,B,C). There was still significant variability again in the TLs between each mutant background, with no particular sets of TEs consistently exhibiting increased transposition (Fig. 3D,E,F).

**Figure 3.**
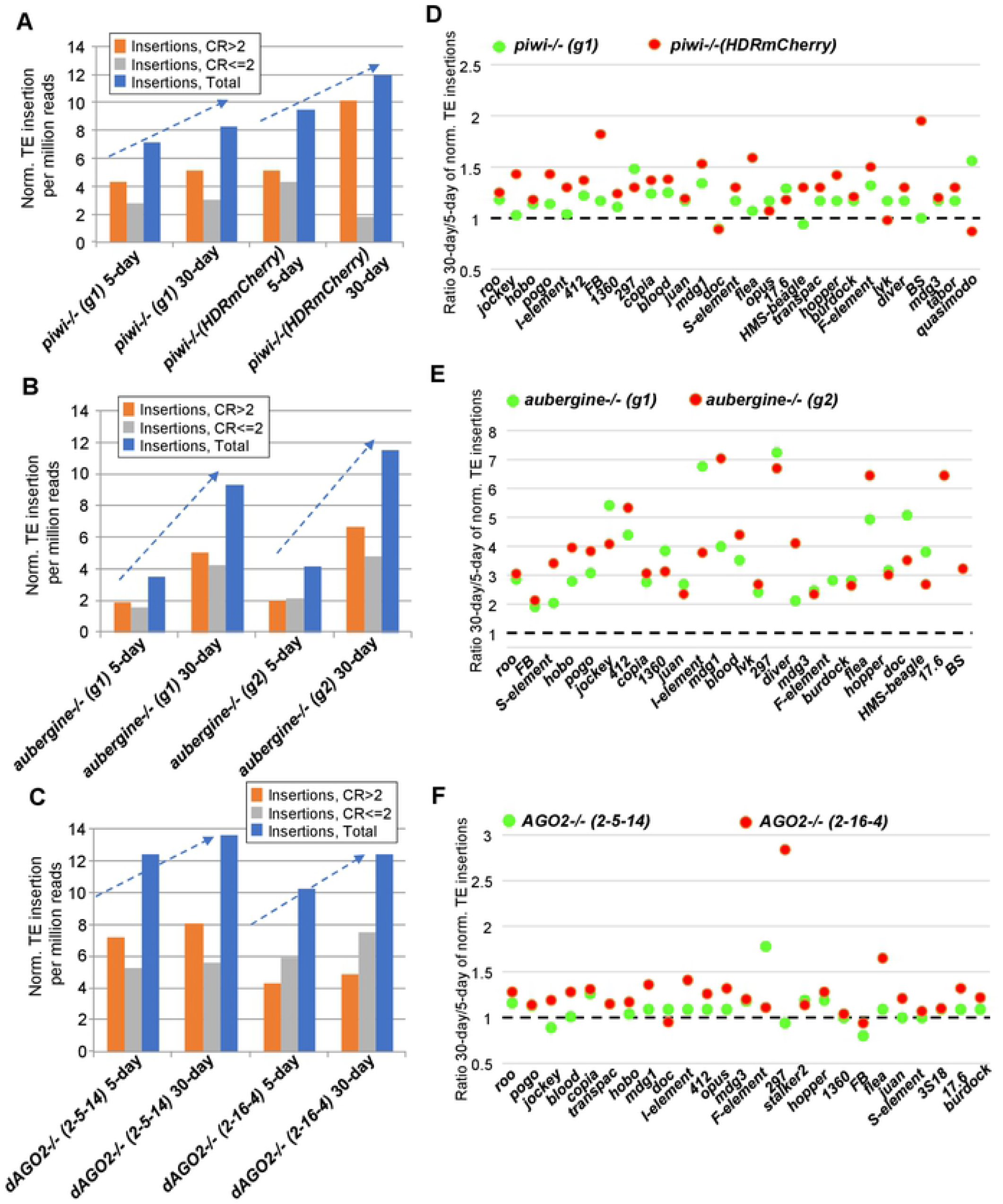
WGS analysis of TE insertion numbers between Young (5-day) versus Aged (30-day) RNAi mutants fly strains. (A-C) Quantification of new TE insertions as compared to the reference genome using the TIDAL-fly program. Categories of total TE insertions broken by the coverage ratios (CR) of CR>2 and CR<=2. Dashed arrows highlight the accumulation of TE insertions in a single generation of aging in two distinct strains of each RNAi null mutants in *piwi-/-, aub-/-, and AGO2*-/- genes. (D-F) Ratios of the 30-day versus 5-day of normalized TE insertions from panels A-C. These panels display only the TE families that were detected by TIDAL be at least 1% of total number of TE families (*i.e*. all the TEs not lumped into the “Others” category of Fig. 2B).

The ddPCR results above affirm that genomic approaches are capable of detecting TL changes and WGS analysis of RNAi mutants demonstrate that the TL changes can be detected during a single generation of aging flies. Therefore, we conclude that although fly aging may allow TE RNAs to become upregulated [10, 13], the RNAi pathways are still functioning in aging WT flies to mitigate TE RNAs from transposing in genomes. In essence, the RNAi mutants raise the frequency of successful TE mobilization events earlier in development so that these new insertions become highly penetrant in the pooled population of the WGS libraries. This would also suggest that successful TE insertions in WT flies are usually infrequent as to explain the modest changes in WT TLs when sequencing genomes from whole flies.

### Detectable TL changes in fly brains during aging

Perhaps new TE insertions may be better detected in specific tissues were cells that are more permanent and not turned over as frequently, such as the brain. For example, in mammalian neurons, the most active TE *LINE-L1* has been implicated in transposing relatively frequently during development to give rise to genomic mosaicism in the brain [45–53]. Given the caveats of having to do prior total DNA amplification from limited gDNA from fly neurons [60], we undertook WGS from at least 50 dissected female brains to provide sufficient nucleic acid for RT-PCR confirmation of neuronal gene expression and WGS of brain DNA (**Figure 4**).

**Figure 4.**
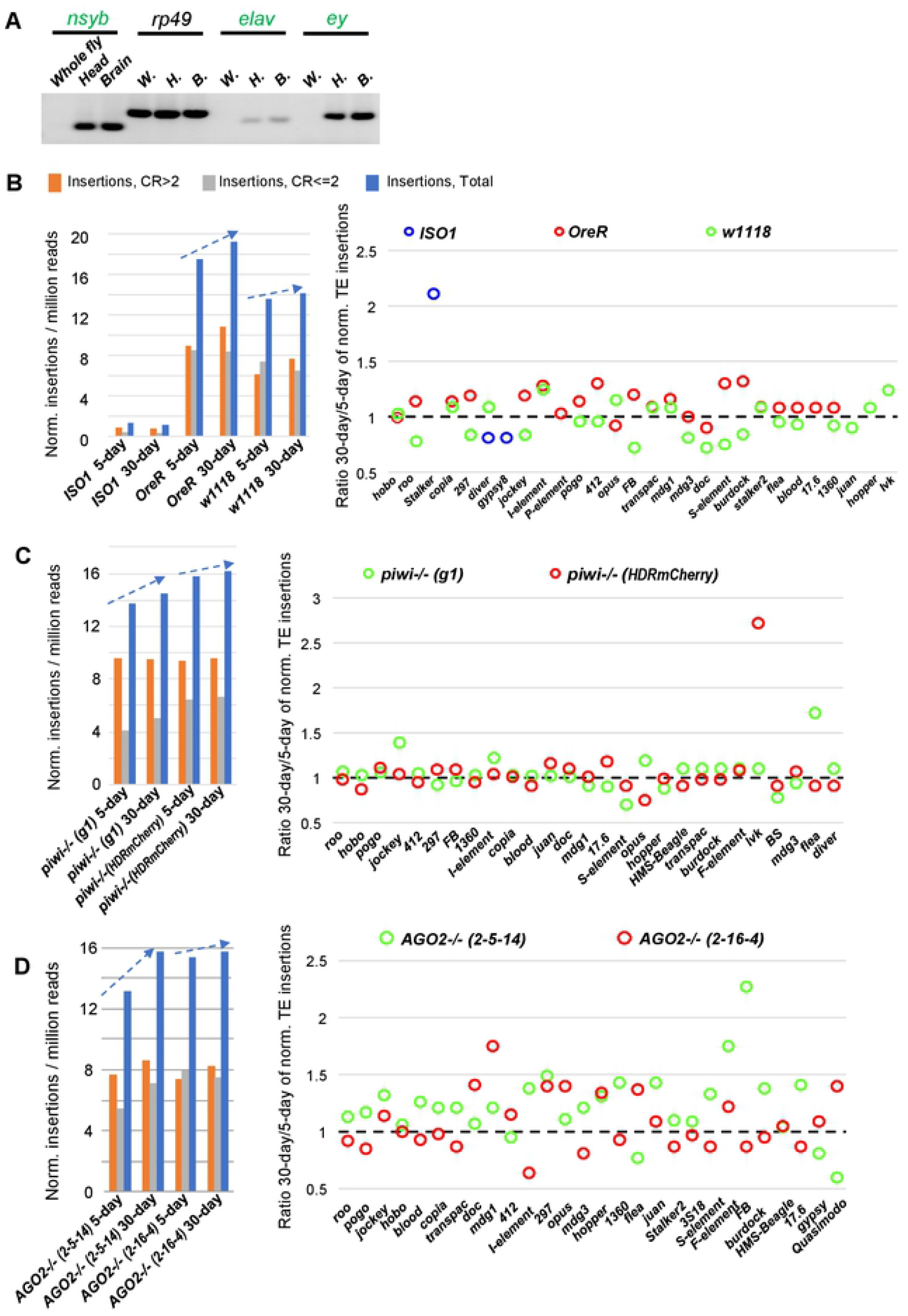
Aging-associated TE landscape changes in fly brains of WT and RNAi mutant strains. (A) Validation of fly brain dissections by microscopy and RT-PCR of brain-specific gene expression. TIDAL analysis of WGS for new TE insertions in the brains of (B) Wild-type (WT) strains, (C) *piwi* mutants, and (D) *Ago2* mutants. The bar graphs on the left represent categories of total TE insertions broken by the Coverage Ratios (CR) of CR>2 and CR<=2. Dashed arrows highlight the accumulation of TE insertions in a single generation of aging flies. The dot graphs to the right show the ratios of the 30-day versus 5-day of normalized TE insertions from left panels B-D. These panels display only the TE families that were detected by TIDAL be at least 1% of total number of TE families, hence only three blue dots for ISO1 are visible in (B).

We successfully generated libraries directly from brains of WT fly strains and *piwi* and *AGO2* mutants without any prior total DNA amplification, and now we could detect increases in TLs from *OreR* and *w1118* strains (Fig. 4B). Although there may be piRNA-like small RNAs and *piwi* expression in fly heads [13, 21, 22], we detected increases in TLs in *piwi* mutants’ brains that were similar in magnitude to the WT *OreR* and *w1118* strains (Fig. 4C). Greater and more variable increases in TLs were apparent in the *AGO2* mutants’ brains (Fig. 4D). Only the brains from the *ISO1* strain were recalcitrant from showing much TE insertion increases except for a >2-fold increase in the *Stalker* TE (Fig. 4B). Except for *ISO1*, the other fly strains appear to enable increasing TL changes in the brain during fly aging.

### Extra-chromosomal circular DNAs (eccDNAs) as an additional genomic cache of increasing TE sequences

In normal and diseased animal cells, there is a cache of eccDNAs that has recently been explored by deep sequencing of DNA that is resistant to extensive exonuclease digestion [81–84]. In certain tumor samples, eccDNAs are implicated in rapid copy-number expansion of oncogenes [85], while ectopic accumulation of DNA in the cytoplasm of senescing cells might trigger aging-associated inflammation responses [86]. Several earlier studies had also found evidence of eccDNAs in *Drosophila*, with the *copia* TE as a prominent example accumulating in certain strains [87–91]. Lastly, eccDNA enriched in TE sequences and other repeats were detected in normal plants and gDNA of human tissues [81, 92], which in both of these studies required total DNA amplification prior to library construction to enrich the surviving eccDNAs after exonuclease digestion.

We investigated eccDNAs in *Drosophila* by optimizing our own method to purify enough eccDNAs to directly generate libraries for deep sequencing without requiring prior total DNA amplification (**Figure 5A**). Furthermore, we used spike-ins of cloning-vector plasmid DNAs into gDNA preparations and magnetic beads for improved recovery and quantitation of eccDNAs for comparing between different samples. To confirm that eccDNA was recovered after two rounds of Exo5 and Exo8 exonuclease digestion steps which only degrade linear but not circular DNA, we conducted PCR with standard primers amplifying linear genes and TEs (F1-R1 primer pairs, **Table S2**), and outward-facing primers that either generate an amplicon from a TE eccDNA or tandem genomic copies of the same TE (P10-P11 primer pairs) (Fig. 5B). Linear gene amplicons were significantly depleted after exonuclease digestions, while the amplicon for the spike-in plasmid was enriched. Linear TE amplicons were also reduced while eccDNA-targeted amplicons for the *copia* TE was resilient against the exonuclease treatment. Some other TE amplicons with outward-facing primers that were reduced after exonuclease treatment may reflect more tandem copies of these TEs.

**Figure 5.**
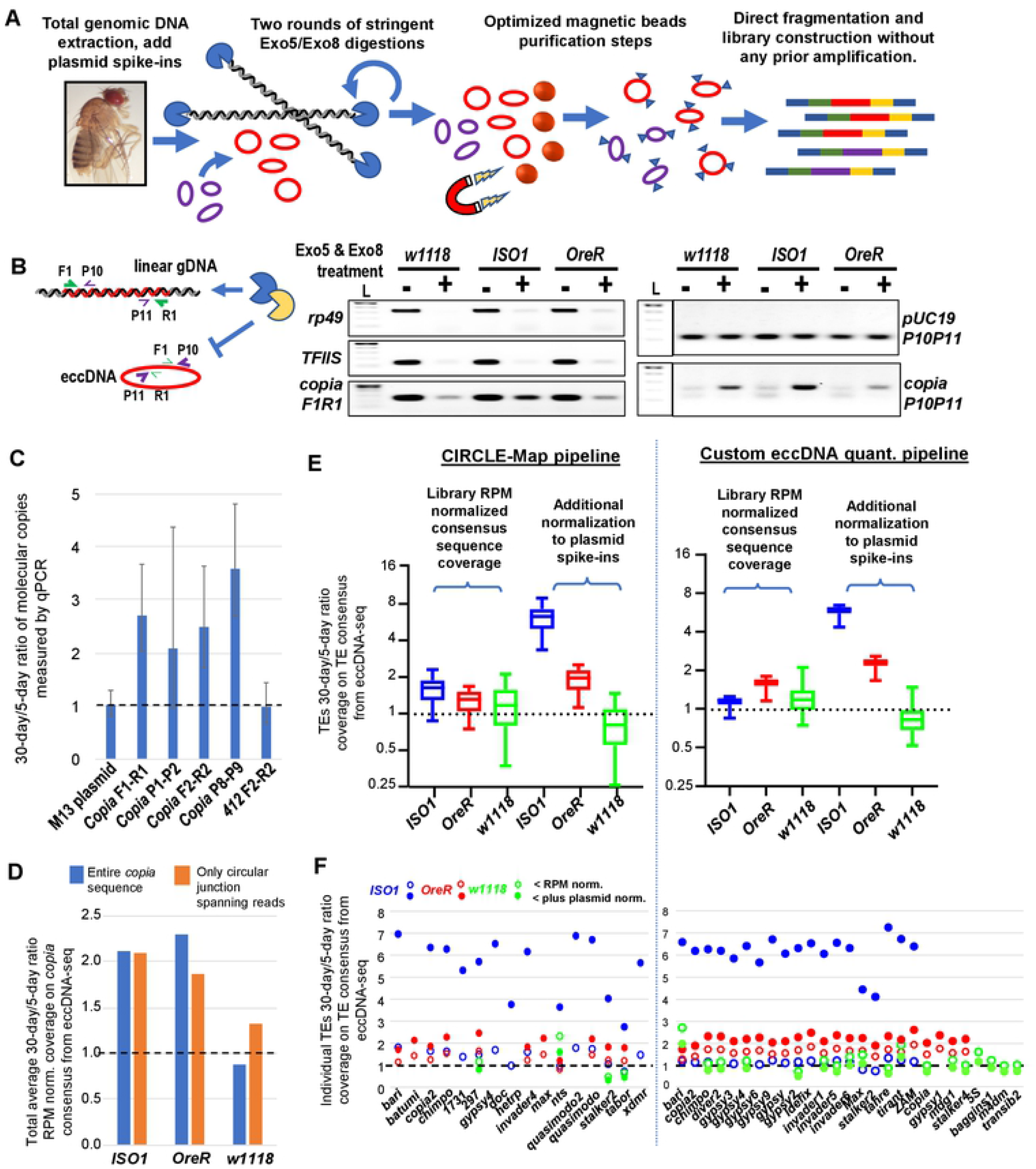
Aging *Drosophila* display increases in TEs existing as extra-chromosomal circular DNA (eccDNA). (A) Diagram of methodology to enrich and purify eccDNAs for direct library construction and sequencing without requiring prior amplification. (B) Genomic PCR from WT flies demonstrating the depletion of linear gDNA and enrichment of eccDNA with TE sequences during exonucleases treatments. Inset diagram explains configuration of PCR primers. Left diagram explains configuration of PCR primers, L=DNA ladder. (C) qPCR validation of spike-in plasmids and *copia* eccDNA after exonucleases treatments of *ISO1* gDNA from young versus aged adult flies. (D) Ratio of the read coverage just across the *copia* consensus sequence comparing young versus aged flies. (E) Box plots of 30-day/5-day ratios of read coverage for eccDNA TE sequences rated by the CIRCLE-Map pipeline with significant “circle score” >50 (Moller et al, 2018); and for our own custom quantitation pipeline that uses a TE-mapping scripts previously used for small RNA analysis. (F) Dot graphs highlighting specific TE eccDNAs whose 30-day/5-day sequencing ratios are normalized to the RPM library size or further normalized to the plasmid spike-ins from (E). These panels display only the TE families that had “circle score” >50 (left) or displayed a measurable 30-day/5-day ratio from the custom analysis pipeline (right).

Since the regular PCR amplicons for the *copia* eccDNA were readily apparent in WT strains (Fig. 5B), we used qPCR to quantify the changes and show that *copia* eccDNA copies were increased >~2-fold in 30-day aged flies compared to 5-day young flies (Fig. 5C). This result motivated us to deeply sequence short read libraries generated directly from those eccDNA-enriched samples which did not undergo any total DNA amplification (**Table S3**). We first adapted the TIDAL scripts of mapping reads to the TE families consensus sequences to measure sequencing coverage as well as circular junction spanning reads against *copia* and observed an aging-associated increase in *copia* eccDNA that was consistent with our qPCR results (Fig. 5D). We also applied this custom eccDNA quantitation pipeline to all the other *Drosophila* TEs as well as adapting the CIRCLE-Map pipeline previously used to measure mammalian eccDNAs [81] to the *Drosophila* TEs. We then normalized the ratios of the eccDNA-TE counts between 30-day aged and 5-day young flies (Fig. 5E). Although the CIRCLE-Map pipeline was more sophisticated at providing a significance “circle score” that we set the cutoff to be >50, our custom eccDNA quantitation pipeline’s results were notably consistent in showing overall that most eccDNAs as TEs were increasing in the libraries of 30-day aged flies (Fig, 5E, F). However, the additional normalization to the plasmid spike-ins were more informative in moderating eccDNA levels in *w1118* while reaffirming the TE eccDNA increases in *OreR* and *ISO1* (Fig, 5E, F). Thus, while *ISO1* TLs did not change much at the chromosomal level during aging, *ISO1* TE copy numbers may instead increase through eccDNA accumulation.

### Genetically enhancing RNAi counteracts TE expression during Drosophila aging

Although TE expression still increased in WT aging flies, we hypothesized whether endogenous RNAi pathways that still limit genomic TL increases could also be genetically enhanced to mitigate the aging-associated rise of TE RNAs. To test this hypothesis, we first used a ubiquitous *Tubulin-GAL4* driver to overexpress *AGO2* in adults, and as expected, multiple TE RNAs had lowered expression relative to the negative control (**Figure 6A**). We then used the same driver to overexpress *piwi*, and although there was likely a silencing limit to prevalent *piwi* expression in the ovary, the enhancement of *piwi* expression and TE silencing was much more apparent in the female carcass (Fig. 6B).

**Figure 6.**
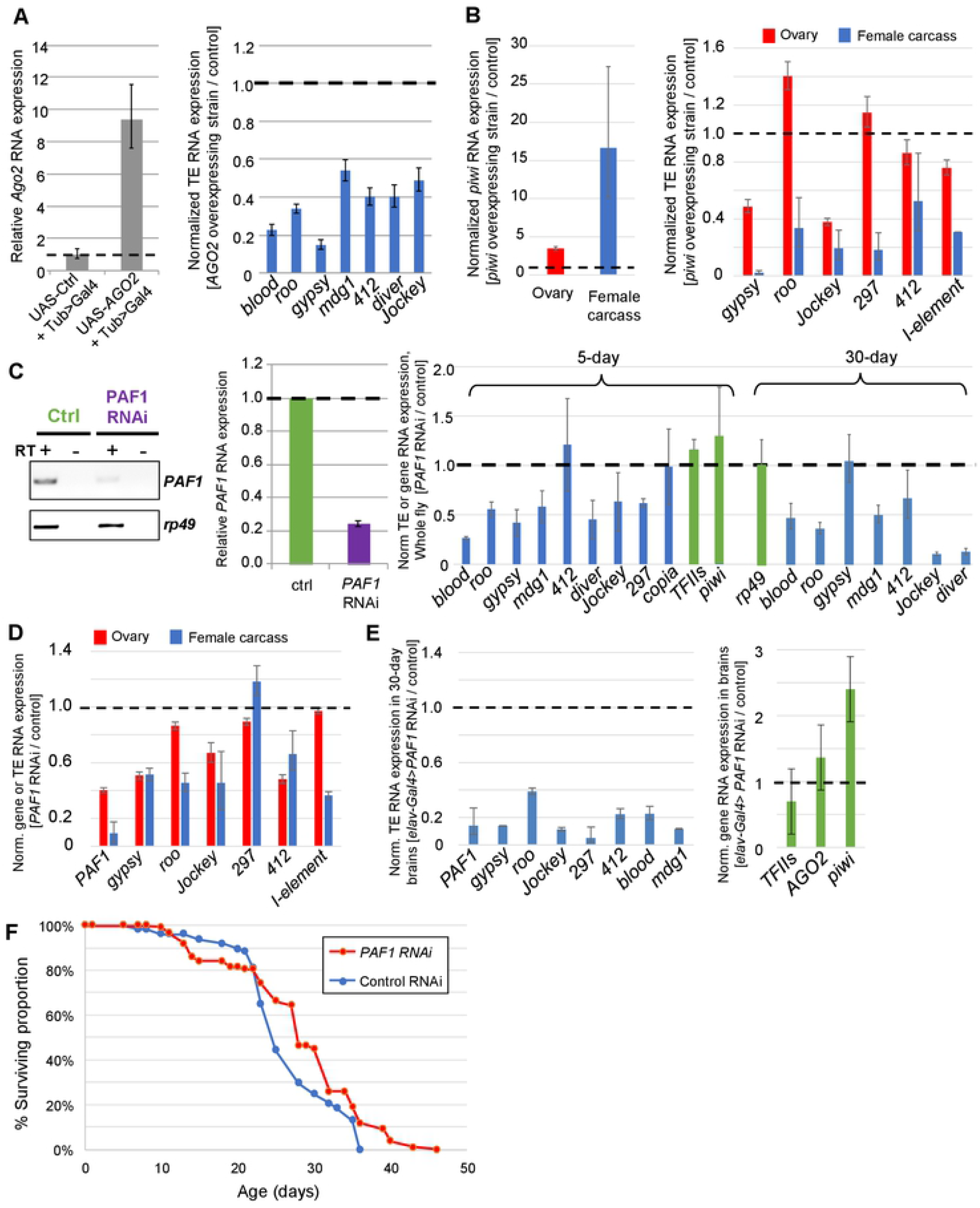
Genetic interventions of TE expression in adult *Drosophila*. (A) Overexpressing AGO2 [Tub>Gal4;UAS-HA-*AGO2*]/[Tub>Gal4] and (B) overexpressing PIWI [Tub>Gal4;UAS-3X-HA *piwi*]/[Tub>Gal4] results in a reduction of TE RNA expression in 5-day young adult *Drosophila*. Left graphs confirm gene overexpression and right graphs detail TE RNA expression measured by RT-qPCR of the target gene compared to the *rp49* housekeeping gene and with error bars representing propagated standard error of triplicate measurements. (C) Adult-specific knockdown of *PAF1* in 5-day young females qualitatively assessed in the gel (left) and RT-qPCR (middle), which reduces TE RNA expression (right). The genes *TFIIs* and *piwi* are controls suggesting that TE RNA reduction is distinct from a concern that *PAF1* RNAi would simply be causing global reduction in transcription. Examining the effect of TE RNA reduction in the *PAF1* knockdown in the ovary (D) and brain (E) of adult *Drosophila* with TEs and *PAF1* in left graph and control genes in the right graph. (F) Life span comparison between control versus *PAF1* RNAi knockdown of adult female flies upon raising them at 29 °C to release the GAL80^ts^ inhibitor to induce RNAi from the Tub>GAL4. *PAF1 RNAi* n=112, Control RNAi n=170.

These data provided a proof of principal that augmenting these RNAi pathways in adults results in improvements in TE silencing. However, inhibiting a factor that normally limits RNAi activity would be preferable from a therapeutic standpoint. Examples of endogenous negative regulation of RNAi activity include proteasome-mediated turnover of *AGO2* [93], ENRI factors that negatively regulate nuclear RNAi in nematodes [94], and the RNA exosome and *PAF1’s* transcription elongation role modulating RNAi silencing activity on TEs conserved in both fission yeast and flies [95–97]. Even though we were able to use siRNA knockdown of *PAF1* in *Drosophila OSS* cells to demonstrate enhanced TE silencing, we recognized that genetic knockdowns of this essential modulator of RNAi would have detrimental effects on development like its requirement in ovarian development [95].

So, to circumvent developmental impacts of *PAF1* knockdown in flies, we further combined the temperature-sensitive inhibitor of GAL4 expressed from a second transgene of *Tubulin-Gal80^ts^* with the *Tubulin-Gal4* driver [98]. This double-transgenic fly could then be crossed to the same UAS-*PAF1*-RNAi line so that flies can develop fully at the permissive temperature of 18 °C, and after eclosion be raised at 29 °C to trigger the RNAi knockdown of *PAF1* (Fig. 6C). Because elevated temperature itself can affect TE silencing activity in flies [99–102], we used an *mCherry-shRNA* strain as a negative control that was also raised at 29 °C at the same time as the *PAF1* knockdowns. There was appreciable enhancement of TE silencing in the whole female flies at both 5-day young and 30-day aged flies (Fig. 6C) with similar levels of TE silencing enhancement between the ovaries and the soma (Fig. 6D). We attribute the increased TE silencing during *PAF1* knockdown to the reduced elongation rate of TE transcripts so that RNAi factors can better engage [95] and not from a global transcription reduction because steady state levels of control gene, *TFIIs, AGO2* and *piwi* were not reduced by *PAF1* knockdown (Fig. 6C,E).

Since we had observed TE landscape activity in the adult fly brain (Fig. 4), we also tested a brain-specific driver, *elav-GAL4*, that was effective at triggering PAF1 knockdown and enhancing TE silencing in the 30-day aged fly brains (Fig. 6E). However, this *elav-GAL4* driver that was likely reducing PAF1 levels in all neurons during embryonic and adult development [103] also presented some problems like reduced lifespan relative to the control, and we have not yet been able to recombine *Tubulin-Gal80ts* with the *elav-GAL4* needed for the post-eclosion knockdown experiment. Thus, we report the measured lifespans from the *PAF1* knockdown versus the mCherry-shRNA negative control with the *Tubulin-Gal80^ts^* and *Tubulin-Gal4* driver cross at 29 °C (Fig. 6F). After an initial dip at 2 weeks, the *PAF1* RNAi knockdown flies ended up living longer than the control and suggested that future pharmacological inhibition of *PAF1* activity in maturing adult animals may be a relevant avenue of intervening with the aging-associated increase in TE expression.

## DISCUSSION

In this study we conducted an analysis of WGS approaches towards assessing changing TLs during *Drosophila* aging, and we found that TL increases are readily detectable in the genomes of aging RNAi mutants such as *piwi*, *aubergine* and *AGO2*. These mutants are viable although others have shown that they have reduced longevity compared to control strains [10, 13, 16], and our data now confirms that unchecked elevation of TE transcripts can result in quantifiable genomic alterations in a single lifetime of flies. However, it was more difficult to detect new TE insertions amongst the gDNA of WT fly strains: we had to focus the TIDAL analyses on specific TE families mobilizing into uniquely-mapping sequences and also count the coverage on TE family consensus sequences (Fig. S3). After showing that an orthogonal quantitation method like ddPCR is consistent with TIDAL’s quantitation of TE copy numbers from WGS of *P-elements* and *I-elements* (Fig. S4), our parsimonious conclusion is that despite aging-associated increases in TE expression during fly aging, the RNAi pathway still protects the fly genomes from massively accumulating new TE insertions.

Despite the compactness and completeness of the *D. melanogaster* genome sequence, technical challenges still remain in fully optimizing WGS approaches to quantify TLs. For example, all current metazoan genome assemblies still suffer from large sequencing gaps in telomeric, centromeric and other repetitive regions that remain unanalyzable. Meanwhile, long-read sequencing like Nanopore and PacBio that could close these gaps are still less economical and not as accurate as the Illumina sequencing platform [104], yet library construction methods for the Illumina platform require sufficient input material for reproducible generation of sequencing libraries. Single-cell WGS is not yet robust enough nor has total DNA amplification approaches been demonstrated to be unhampered by molecule bias, so our study required pools of genomes and non-amplified input DNA samples to reduce the prior concerns. Our study also adds a second dimension to WGS of TLs by incorporating eccDNA as an *in vivo* cache of accumulating TE DNA sequences (Fig. 5). Intriguingly, the *ISO1* strain showed the least chromosomal TL changes yet exhibited the greatest increase in TE-eccDNAs in the whole flies, while the *OreR* and *w1118* strains also showed evidence of TE-eccDNAs accumulating in the brain (**Figure S5A**).

In addition to variations in TLs between WT strains, we also observed differences in TLs between other RNAi mutants that we cannot fully explain. For example, we examined aging-associated TLs from two EMS-induced point mutants of *Dcr-2* (*L811fsx*) and *Dcr-2* (*R416X*) from [105]), the nuclease acting upstream of *AGO2* to generate the siRNAs from TE dsRNAs. However, there was inconsistent and contrary TL differences between young and aged *Drosophila* in these *Dcr-2* mutants whole flies and brains (Fig. S5B, C) as well as in *AGO3* mutants (Fig. S5D, [106]). Perhaps these sets of mutants are not as penetrant in the loss of RNAi activity as the *piwi, aubergine and AGO2* mutants. Furthermore, the analysis of a partially rescuing *AGO2* transgene in the *AGO2* (*2-5-14*) null mutant did lower the initial levels of TE insertion differences noted by TIDAL, but the partial rescue still did not fully prevent aging-associated TE increases (Fig. S5E), suggesting only wild type strength RNAi can buffer aging genomes from accumulating new TE insertions.

Therefore, we propose that RNAi activity must be sustained during aging to mitigate negative effects of increased TE expression in aged flies, a phenotype that has also been frequently observed in mammals [34, 55, 107, 108]. To combat TEs’ impact on aging, some therapeutic approaches have used reverse transcriptase inhibitors and drugs that inhibit *LINE-L1* activity [33], while other studies showed that dietary restriction and prolonged exercise in animals can reduce aging-associated increases in TE expression [14, 54, 109]. Our study proposes an additional therapeutic target of augmenting the RNAi pathway’s response to TEs by inhibiting *PAF1*, which has a conserved impact on limiting RNAi from silencing TE transcripts [95, 96]. Perhaps therapeutic siRNAs against *PAF1* transcripts can be hypothesized as a feed-forwarding therapeutic agent to augment RNAi activity in aging animal cells.

A final question to resolve in the future is what cascade of epigenetic and chromatin landscape changes during animal aging consistently leads to increases in TE expression? Given the pleiotropic nature of the animal aging process, we anticipate that there will also be multiple genomic mechanisms that will vary in impact between different genetic backgrounds. For example, we describe variation amongst three WT *Drosophila* strains in the level of accumulating eccDNAs containing TE sequences (Fig. 5), while others have shown increased in polyploidy in adult *Drosophila* brains [110] as well as somatic genome instability in regions of the *Drosophila* genome [111] that might contribute to changes at the level of TE consensus sequence coverages (Fig. S3B). Lastly, during fly aging there are also gross-level changes in histone marks typically associated with chromatin silencing [12, 14], which may precede the increase TE expression, so the future extension of this work will be to add epigenetic and chromatin accessibility landscapes to TLs during *Drosophila* aging.

## ACKNOWLEDGEMENTS

We thank Michael Marr, Dianne Schwarz and Michael Blower for comments on the manuscript. NY, SPS, MC, conducted the experiments; RR, QM developed and executed the bulk of the bioinformatics analyses, GD also provided extensive help with sequencing uploading. MR and EL provided supervisory oversight of RR and MC, respectively. NCL conceived and conducted data analysis and wrote the paper with comments from all authors. This work was supported by NIH grants R01-AG052465 to NCL and a subcontract to MR, and R21-HD088792 to NCL; and Intramural Program of the National Institute of Diabetes and Digestive and Kidney Diseases, National Institutes of Health (DK015602 to E.P.L.). All genomic sequencing data is deposited in the NCBI SRA as Study Bioproject PRJNA678741.

## MATERIALS AND METHODS

### *Drosophila* strains, genetic crosses and aging curves

All flies were raised at 25 °C on standard cornmeal food. For fly aging analyses, newly eclosed female flies were harvested from bottles and mated with males for two days. These females were then divided into ~20 individuals per vial and flipped to new vials every 2-3 days to mitigate crowding stress according to this protocol [112]. Surviving flies were counted at each flip, and the percentage of cumulative survival rate at each time point was plotted against its corresponding age (date of counting subtracted by date of eclosion).

The isogenized *ISO1* fly strain for the Dm6 reference genome sequence was obtained from Susan Celniker [4]; the *w1118* is an isogenized strain and was a gift from R. Scott Hawley [68]; and the *OreR* from the ModEncode project was a gift from Terry Orr-Weaver [69]. The RNAi pathway null mutant strains *piwi-(g1), aubergine-(g1), aubergine-(g2), AGO3-(g1)* and *AGO3-(g2)* were a gift from Julius Brennecke [106]. An additional mutant strain of *Piwi-[HDR-4xP3-mCherry]* [113] was a gift from Eric Lai. The null *AGO2* mutants deletion strains of *AGO2-[2-5-14]* and *AGO2-[2-16-4]* and Ago2-WT-rescue stocks were generated by CRISPR Cas9 approaches as described in [114, 115]. The strains with active and inactive *I-elements* and spermless males were a gift from Zhao Zhang [74]. The UAS-Ago2-HA strain was a gift from Arno Muller lab [116] and the UASp-3xHA-Piwi was a gift from the Ruth Lehman lab [117]. The driver strains of *Tubulin-Gal4* and *elav*-Gal4 were a gift from Leslie Griffith [118]. In addition to a *Tubulin-Gal80^ts^* strain we received from the Griffith lab, we also obtained a second *Tubulin-Gal80^ts^* stock from the Bloomington *Drosophila* Stock Center (BDSC#7018) and combined with *Tubulin-Gal4* for further experiments. The *PAF1* knockdown RNAi line was obtained from the Vienna *Drosophila* Resource Center (VDRC#108826) and an mCherry shRNA control line was obtained from the Harvard TRiP resource (BDSC#35785).

To quantify *I-element* copies by ddPCR, fly cross schemes from [74] were replicated (Fig. S4C). Parental crosses between the *w^1118^* strain with active *I-elements* and *w^k^* strain with inactive *I-elements* were performed reciprocally to generate many virgin F1 females where one strain enables I-element transposition (“invaded” from *w^k^* as the maternal parent) versus a control that maintains I-element silencing (*w^1118^* as the maternal parent). These F1 females were then crossed to sperm-less males that were obtained as F1 male progenies from the parental cross of *w^1118^* virgin females with XY attached male. F2 oocytes were collected overnight and DNA was extracted for ddPCR against the *I-element* and *Rp49*.

### Fly brain isolation, genomic DNA extraction, WGS library construction and deep sequencing

Fly brains were dissected from at least 50 females per age group, following a procedure laid out in [119]. Eye disks and other tissues were removed from heads with forceps, and brain lobes were dissected into tubes with ice-cold PBS before freezing once at −20 °C. Whole female flies and fly brains were homogenized in a standard DNA digestion buffer (1% SDS, 50 mM Tris-HCl pH 8.0, 100 mM EDTA, 100 mM NaCl, 0.5mg/ml Proteinase K) overnight at 50 °C, and then extracted using standard phenol chloroform extraction, ethanol precipitation, and resuspending gDNA pellets in pure water.

WGS of whole flies began with the circa 2014 Nextera Tn5 tagmentation kit (Illumina) using an input of 50 ng gDNA and outputs were purified with AMpure XP beads (Beckman Coulter). WGS libraries were quality controlled with the high-sensitivity DNA kit on the Bioanalyzer (Agilent), selecting for size distributions of 300bp to 1kb and concentrations over 1 nM. Multiplexed libraries were sequenced on Illumina Nextseq500 high-output flow cells using 75 bp paired-end and single end kits. All WGS libraries were sequenced to a minimum depth of 35 million reads (Table S1, S2). After determining that some whole fly libraries made using NEBNext Ultra-II DNA library prep kit for Illumina (NEB) were as complete and has better yields than the then discontinued Nextera kit, we completed the fly brain gDNA libraries with the NEBNext kit and sequenced them to similar depths as above.

### RNA extraction, quantitative RT-PCR, digital droplet PCR (dd-PCR) and TE copy number estimation

Total RNA was extracted from 5-10 female flies harvested at corresponding age with TRI-reagent (MRC, Inc.). Reverse transcription (RT) was performed using random primers, ProtoScript II (NEB), and 1 μg input of total RNA. Quantitative PCR (qPCR) with the Luna Sybr-Green mastermix (NEB) used primer sequences in Table S3 and 2 μL of a 1:10 dilution of the cDNA. Relative changes in gene expression were calculated using the 2^ΔΔCt method with *Rp49* as a housekeeping gene for normalization.

Droplet digital PCR (ddPCR) was conducted on a QX200 instrument with the Evagreen assay reaction (Biorad). Copy number measurements from specific TE primers (Table S3) were normalized to *Rp49* as a diploid gene, starting first at 2 ng of gDNA as input per 20 μL ddPCR for droplet generation for most TEs. For TEs with very high copy numbers that saturate the droplets, input gDNA was diluted further to 2 ng into the ddPCR mix prior to droplet generation. At least 10,000 droplets were required to achieve good statistical estimation of the concentration calculated by Poisson distribution using Quantasoft Analysis Pro (Biorad). TE copy numbers per genome was determined by dividing against half of the measured *Rp49* copies.

### Extracellular circular DNA isolation and sequencing

To quantify eccDNAs during fly aging, 30 female flies were harvested from 5-days and 30-days post eclosion, and a fixed amount off pre-extraction plasmids was added prior to cell lysis: ~80 pg of ~7kb-pGL3-DmPiwipro1 and ~50 pg of ~11kb-pCas9 prior to cell lysis. About 30 ug of total gDNA was recovered from using MasterPure™ Complete DNA and RNA Purification kit (Lucigen), and 0.5ug-1ug gDNA was checked on a 1% agarose gel for integrity and quality. Good gDNA primarily migrated at >10kb and to 20 μg gDNA we added 40ul of a second plasmid cocktail: (1ng/ul of the 2.7kb pUC19, 0.1ng/ul of the 3.5kb pMaxGFP, 0.01ng/ul of the 5.2kb pGSH0 and 0.001ng/ul of the 6.3kb pCENPm3) and split equally to two reactions: Exo5/8 non-treated control versus Exo5/8 treated samples. We conducted a first round of Exo5/Exo8 (NEB) treatment at 37 °C overnight, then an additional 2-hour treatment with freshly replenished buffer, ATP and enzymes. The reaction was stopped and purified using AMPure XP beads (Beckman Coulter) and eluted in 50 ul of water.

To check the efficiency of Exo5/8 treatment, 10 ul of the eluate from untreated versus treated samples were loaded on 1% agarose gel to visualize complete digestion of gDNA. We quality controlled Exo5/8 treatments by performing qPCR against rp49, ND5 and various plasmid primers including pUC19, pGL3piwipro and pCas9 and Ct values were compared between untreated versus treated samples. Mitochondria was not a reliable circular molecule because of the high variability of ND5 Ct values across multiple sample preps. Comparing between treated and untreated sample, the plasmids Ct values were generally stable (<2 Ct difference), and much higher for rp49 (>5 Ct difference) indicating the Exo5/8 treatments were effective at removing linear chromosomal DNA and not affecting the circular plasmids. Half of the Exo5/8 treated sample (25 ul out of 50 ul purified elute) was used as template for library construction using NEBNext UltraII library prep kit as stated above. Libraries were single end (75bp) or pair-end sequenced at 36 bp by 36 bp on a Nextseq550 flow cell (Illumina).

For eccDNA sequencing from brains, 200 female brains were dissected and added with half the volume of pre-extraction plasmids as whole flies, and gDNA concentration was measured by the Qubit 4 Fluorometer (Thermofisher). To 100 ng of brain gDNA, we mixed 20 ul of the plasmid spike-in cocktail and a tenth of the Exo5/8 enzyme as whole flies gDNA. At least 10 million eccDNA reads were required for analysis.

### TIDAL updates with total TE consensus and gene mapping strategies

TE insertion analysis was carried out with an updated version of our previously developed TIDAL program (original code available on the Github repository at: https://github.com/laulabbrandeis/TIDAL) [65]. In this study, the updated version of TIDALv1.2 is also posted to Github at (https://github.com/laulabbumc/TIDAL1.2). These scripts carry out the analysis run the same way as the original TIDAL, but we incorporated two additional features. First, for the euchromatic TE insertions we selected 22 arbitrarily selected protein coding gene (Immobile gene elements IGEs) that are computed along with consensus TE sequence to benchmark noise in detection of genetic elements. The algorithm used to identify transposon insertion sites based on consensus transposon sequence is then applied on these 100 IGE sequence to determine their insertion sites. Second, for the total reads mapped to consensus TE sequences, here we added 100 IGEs are computed by mapping reads with bowtie2 using parameters “--sensitive --end-to-end” and custom shell, Perl, C-code, and R-code scripts all accessible from (https://github.com/laulabbumc/TIDAL1.2).

TEMP v1.05 code was acquired from the GitHub repository at: (https://github.com/JialiUMassWengLab/TEMP), and was run with default parameters except “-x 30, -m 3 -f 500”. These parameters were chosen to ensure that TEMP results are consistent with analysis shown in [60, 62].

### Bioinformatics counting of eccDNA from TEs and spike-in plasmids using a custom pipeline and CIRCLE-Map program

In our first look at the eccDNA reads, we inputted them into an existing bioinformatics pipeline already developed for mapping *Drosophila* small RNA counts to TEs [120]. Reads were first checked by the Cutadapt program to see if adaptor sequences at the 3’ end needed to be removed, and then we indexed the reads to the *Drosophila* genome assembly file by running BWA version 1 [121] and formatdb from NCBI. Using Bowtie1 with 2 mismatches [122], reads were mapped to genome to get the genic and intergenic counts using the genome GTF file. The total number of reads mapped to the *Drosophila* genome was derived by subtracting the total number of reads not mapped to the *Drosophila* genome from the total number of reads. The total number of mapped reads was used as the basis for normalization of TE counts and spike-in plasmid counts.

Plasmid sequences were treated as linear entries in the FASTA file database similar to the TE family consensus sequences. The raw read counts from TE mapping were further normalized by the total number of reads mapping to the *Drosophila* Dm6 genome assembly. For spike-in plasmid counting, because several plasmids share the same backbone with different inserts, read frequencies were normalized by the total plasmid mapping sites as well as by the total number of *Drosophila* genome-mapping reads.

To execute the CIRCLE-Map program for repeats [81], we indexed the *Drosophila* genome FASTA file by BWA. We then used the MEM algorithm under BWA to align reads against the *Drosophila* genome FASTA file. Next, we sorted the reads by alignment position within the resulting BAM file and indexed the resulting BAM file. Finally, we detected the circles by calling CIRCLE-Map program. The CIRCLE-Map program for repeats yields an output for reads with two high scoring alignments as these ones are indicative of circles formed from regions with homology.

## SUPPORTING ONLINE MATERIALS LIST

**Supplementary Text, Supplementary Figures and Tables legends.**

**Figures S1-S5.**

**Tables S1-S3.**

## REFERENCES

1. Smith JJ, Timoshevskaya N, Timoshevskiy VA, Keinath MC, Hardy D, Voss SR. A chromosome-scale assembly of the axolotl genome. Genome research. 2019;29(2):317–24. Epub 2019/01/27. doi: 10.1101/gr.241901.118. PubMed PMID: 30679309; PubMed Central PMCID: PMCPMC6360810.

2. Nowoshilow S, Schloissnig S, Fei JF, Dahl A, Pang AWC, Pippel M, et al. The axolotl genome and the evolution of key tissue formation regulators. Nature. 2018;554(7690):50–5. Epub 2018/01/25. doi: 10.1038/nature25458. PubMed PMID: 29364872.

3. Lander ES, Linton LM, Birren B, Nusbaum C, Zody MC, Baldwin J, et al. Initial sequencing and analysis of the human genome. Nature. 2001;409(6822):860–921. doi: 10.1038/35057062. PubMed PMID: 11237011.

4. Adams MD, Celniker SE, Holt RA, Evans CA, Gocayne JD, Amanatides PG, et al. The genome sequence of Drosophila melanogaster. Science. 2000;287(5461):2185–95. PubMed PMID: 10731132.

5. Kaminker JS, Bergman CM, Kronmiller B, Carlson J, Svirskas R, Patel S, et al. The transposable elements of the Drosophila melanogaster euchromatin: a genomics perspective. Genome biology. 2002;3(12):RESEARCH0084. PubMed PMID: 12537573; PubMed Central PMCID: PMC151186.

6. Chuong EB, Elde NC, Feschotte C. Regulatory activities of transposable elements: from conflicts to benefits. Nature reviews Genetics. 2017;18(2):71–86. doi: 10.1038/nrg.2016.139. PubMed PMID: 27867194; PubMed Central PMCID: PMCPMC5498291.

7. Wells JN, Feschotte C. A Field Guide to Eukaryotic Transposable Elements. Annual review of genetics. 2020. Epub 2020/09/22. doi: 10.1146/annurev-genet-040620-022145. PubMed PMID: 32955944.

8. Gorbunova V, Boeke JD, Helfand SL, Sedivy JM. Human Genomics. Sleeping dogs of the genome. Science. 2014;346(6214):1187–8. doi: 10.1126/science.aaa3177. PubMed PMID: 25477445; PubMed Central PMCID: PMC4312280.

9. Chen H, Zheng X, Xiao D, Zheng Y. Age-associated de-repression of retrotransposons in the Drosophila fat body, its potential cause and consequence. Aging Cell. 2016;15(3):542–52. Epub 2016/04/14. doi: 10.1111/acel.12465. PubMed PMID: 27072046; PubMed Central PMCID: PMCPMC4854910.

10. Li W, Prazak L, Chatterjee N, Gruninger S, Krug L, Theodorou D, et al. Activation of transposable elements during aging and neuronal decline in Drosophila. Nature neuroscience. 2013;16(5):529–31. doi: 10.1038/nn.3368. PubMed PMID: 23563579; PubMed Central PMCID: PMC3821974.

11. Wood JG, Helfand SL. Chromatin structure and transposable elements in organismal aging. Frontiers in genetics. 2013;4:274. doi: 10.3389/fgene.2013.00274. PubMed PMID: 24363663; PubMed Central PMCID: PMC3849598.

12. Wood JG, Hillenmeyer S, Lawrence C, Chang C, Hosier S, Lightfoot W, et al. Chromatin remodeling in the aging genome of Drosophila. Aging Cell. 2010;9(6):971–8. doi: 10.1111/j.1474-9726.2010.00624.x. PubMed PMID: 20961390; PubMed Central PMCID: PMC2980570.

13. Jones BC, Wood JG, Chang C, Tam AD, Franklin MJ, Siegel ER, et al. A somatic piRNA pathway in the Drosophila fat body ensures metabolic homeostasis and normal lifespan. Nature communications. 2016;7:13856. doi: 10.1038/ncomms13856. PubMed PMID: 28000665; PubMed Central PMCID: PMCPMC5187580.

14. Wood JG, Jones BC, Jiang N, Chang C, Hosier S, Wickremesinghe P, et al. Chromatin-modifying genetic interventions suppress age-associated transposable element activation and extend life span in Drosophila. Proceedings of the National Academy of Sciences of the United States of America. 2016;113(40):11277–82. doi: 10.1073/pnas.1604621113. PubMed PMID: 27621458; PubMed Central PMCID: PMCPMC5056045.

15. Chang YH, Keegan RM, Prazak L, Dubnau J. Cellular labeling of endogenous retrovirus replication (CLEVR) reveals de novo insertions of the gypsy retrotransposable element in cell culture and in both neurons and glial cells of aging fruit flies. PLoS biology. 2019;17(5):e3000278. Epub 2019/05/17. doi: 10.1371/journal.pbio.3000278. PubMed PMID: 31095565; PubMed Central PMCID: PMCPMC6541305.

16. Sousa-Victor P, Ayyaz A, Hayashi R, Qi Y, Madden DT, Lunyak VV, et al. Piwi Is Required to Limit Exhaustion of Aging Somatic Stem Cells. Cell reports. 2017;20(11):2527–37. Epub 2017/09/14. doi: 10.1016/j.celrep.2017.08.059. PubMed PMID: 28903034; PubMed Central PMCID: PMCPMC5901960.

17. Chang YH, Dubnau J. The Gypsy Endogenous Retrovirus Drives Non-Cell-Autonomous Propagation in a Drosophila TDP-43 Model of Neurodegeneration. Curr Biol. 2019;29(19):3135–52 e4. Epub 2019/09/10. doi: 10.1016/j.cub.2019.07.071. PubMed PMID: 31495585; PubMed Central PMCID: PMCPMC6783360.

18. Krug L, Chatterjee N, Borges-Monroy R, Hearn S, Liao WW, Morrill K, et al. Retrotransposon activation contributes to neurodegeneration in a Drosophila TDP-43 model of ALS. PLoS genetics. 2017;13(3):e1006635. doi: 10.1371/journal.pgen.1006635. PubMed PMID: 28301478; PubMed Central PMCID: PMCPMC5354250.

19. Sun W, Samimi H, Gamez M, Zare H, Frost B. Pathogenic tau-induced piRNA depletion promotes neuronal death through transposable element dysregulation in neurodegenerative tauopathies. Nature neuroscience. 2018;21(8):1038–48. Epub 2018/07/25. doi: 10.1038/s41593-018-0194-1. PubMed PMID: 30038280; PubMed Central PMCID: PMCPMC6095477.

20. Guo C, Jeong HH, Hsieh YC, Klein HU, Bennett DA, De Jager PL, et al. Tau Activates Transposable Elements in Alzheimer’s Disease. Cell reports. 2018;23(10):2874–80. Epub 2018/06/07. doi: 10.1016/j.celrep.2018.05.004. PubMed PMID: 29874575; PubMed Central PMCID: PMCPMC6181645.

21. Mirkovic-Hosle M, Forstemann K. Transposon defense by endo-siRNAs, piRNAs and somatic pilRNAs in Drosophila: contributions of Loqs-PD and R2D2. PloS one. 2014;9(1):e84994. Epub 2014/01/24. doi: 10.1371/journal.pone.0084994. PubMed PMID: 24454776; PubMed Central PMCID: PMCPMC3890300.

22. Ghildiyal M, Seitz H, Horwich MD, Li C, Du T, Lee S, et al. Endogenous siRNAs derived from transposons and mRNAs in Drosophila somatic cells. Science. 2008;320(5879):1077–81. Epub 2008/04/12. doi: 1157396 [pii] 10.1126/science.1157396. PubMed PMID: 18403677.

23. Lin KY, Wang WD, Lin CH, Rastegari E, Su YH, Chang YT, et al. Piwi reduction in the aged niche eliminates germline stem cells via Toll-GSK3 signaling. Nature communications. 2020;11(1):3147. Epub 2020/06/21. doi: 10.1038/s41467-020-16858-6. PubMed PMID: 32561720; PubMed Central PMCID: PMCPMC7305233.

24. Kato M, Takemoto K, Shinkai Y. A somatic role for the histone methyltransferase Setdb1 in endogenous retrovirus silencing. Nature communications. 2018;9(1):1683. Epub 2018/04/29. doi: 10.1038/s41467-018-04132-9. PubMed PMID: 29703894; PubMed Central PMCID: PMCPMC5923290.

25. Matsui T, Leung D, Miyashita H, Maksakova IA, Miyachi H, Kimura H, et al. Proviral silencing in embryonic stem cells requires the histone methyltransferase ESET. Nature. 2010;464(7290):927–31. Epub 2010/02/19. doi: 10.1038/nature08858. PubMed PMID: 20164836.

26. Ecco G, Cassano M, Kauzlaric A, Duc J, Coluccio A, Offner S, et al. Transposable Elements and Their KRAB-ZFP Controllers Regulate Gene Expression in Adult Tissues. Developmental cell. 2016;36(6):611–23. Epub 2016/03/24. doi: 10.1016/j.devcel.2016.02.024. PubMed PMID: 27003935; PubMed Central PMCID: PMCPMC4896391.

27. Rowe HM, Jakobsson J, Mesnard D, Rougemont J, Reynard S, Aktas T, et al. KAP1 controls endogenous retroviruses in embryonic stem cells. Nature. 2010;463(7278):237–40. Epub 2010/01/16. doi: 10.1038/nature08674. PubMed PMID: 20075919.

28. Castro-Diaz N, Ecco G, Coluccio A, Kapopoulou A, Yazdanpanah B, Friedli M, et al. Evolutionally dynamic L1 regulation in embryonic stem cells. Genes & development. 2014;28(13):1397–409. Epub 2014/06/19. doi: 10.1101/gad.241661.114. PubMed PMID: 24939876; PubMed Central PMCID: PMCPMC4083085.

29. Tunbak H, Enriquez-Gasca R, Tie CHC, Gould PA, Mlcochova P, Gupta RK, et al. The HUSH complex is a gatekeeper of type I interferon through epigenetic regulation of LINE-1s. Nature communications. 2020;11(1):5387. Epub 2020/11/05. doi: 10.1038/s41467-020-19170-5. PubMed PMID: 33144593; PubMed Central PMCID: PMCPMC7609715.

30. Robbez-Masson L, Tie CHC, Conde L, Tunbak H, Husovsky C, Tchasovnikarova IA, et al. The HUSH complex cooperates with TRIM28 to repress young retrotransposons and new genes. Genome research. 2018;28(6):836–45. Epub 2018/05/08. doi: 10.1101/gr.228171.117. PubMed PMID: 29728366; PubMed Central PMCID: PMCPMC5991525.

31. Liu N, Lee CH, Swigut T, Grow E, Gu B, Bassik MC, et al. Selective silencing of euchromatic L1s revealed by genome-wide screens for L1 regulators. Nature. 2018;553(7687):228–32. Epub 2017/12/07. doi: 10.1038/nature25179. PubMed PMID: 29211708; PubMed Central PMCID: PMCPMC5774979.

32. Tchasovnikarova IA, Timms RT, Matheson NJ, Wals K, Antrobus R, Gottgens B, et al. GENE SILENCING. Epigenetic silencing by the HUSH complex mediates position-effect variegation in human cells. Science. 2015;348(6242):1481–5. Epub 2015/05/30. doi: 10.1126/science.aaa7227. PubMed PMID: 26022416; PubMed Central PMCID: PMCPMC4487827.

33. Simon M, Van Meter M, Ablaeva J, Ke Z, Gonzalez RS, Taguchi T, et al. LINE1 Derepression in Aged Wild-Type and SIRT6-Deficient Mice Drives Inflammation. Cell metabolism. 2019;29(4):871–85 e5. Epub 2019/03/12. doi: 10.1016/j.cmet.2019.02.014. PubMed PMID: 30853213; PubMed Central PMCID: PMCPMC6449196.

34. Van Meter M, Kashyap M, Rezazadeh S, Geneva AJ, Morello TD, Seluanov A, et al. SIRT6 represses LINE1 retrotransposons by ribosylating KAP1 but this repression fails with stress and age. Nature communications. 2014;5:5011. doi: 10.1038/ncomms6011. PubMed PMID: 25247314; PubMed Central PMCID: PMCPMC4185372.

35. Bulut-Karslioglu A, De La Rosa-Velazquez IA, Ramirez F, Barenboim M, Onishi-Seebacher M, Arand J, et al. Suv39h-dependent H3K9me3 marks intact retrotransposons and silences LINE elements in mouse embryonic stem cells. Molecular cell. 2014;55(2):277–90. Epub 2014/07/02. doi: 10.1016/j.molcel.2014.05.029. PubMed PMID: 24981170.

36. Leung DC, Dong KB, Maksakova IA, Goyal P, Appanah R, Lee S, et al. Lysine methyltransferase G9a is required for de novo DNA methylation and the establishment, but not the maintenance, of proviral silencing. Proceedings of the National Academy of Sciences of the United States of America. 2011;108(14):5718–23. Epub 2011/03/24. doi: 10.1073/pnas.1014660108. PubMed PMID: 21427230; PubMed Central PMCID: PMCPMC3078371.

37. Vogel MJ, Guelen L, de Wit E, Peric-Hupkes D, Loden M, Talhout W, et al. Human heterochromatin proteins form large domains containing KRAB-ZNF genes. Genome research. 2006;16(12):1493–504. Epub 2006/10/14. doi: 10.1101/gr.5391806. PubMed PMID: 17038565; PubMed Central PMCID: PMCPMC1665633.

38. Schopp T, Zoch A, Berrens RV, Auchynnikava T, Kabayama Y, Vasiliauskaite L, et al. TEX15 is an essential executor of MIWI2-directed transposon DNA methylation and silencing. Nature communications. 2020;11(1):3739. Epub 2020/07/29. doi: 10.1038/s41467-020-17372-5. PubMed PMID: 32719317; PubMed Central PMCID: PMCPMC7385494.

39. Zoch A, Auchynnikava T, Berrens RV, Kabayama Y, Schopp T, Heep M, et al. SPOCD1 is an essential executor of piRNA-directed de novo DNA methylation. Nature. 2020;584(7822):635–9. Epub 2020/07/17. doi: 10.1038/s41586-020-2557-5. PubMed PMID: 32674113.

40. Carmell MA, Girard A, van de Kant HJ, Bourc’his D, Bestor TH, de Rooij DG, et al. MIWI2 Is Essential for Spermatogenesis and Repression of Transposons in the Mouse Male Germline. Developmental cell. 2007. PubMed PMID: 17395546.

41. Kuramochi-Miyagawa S, Watanabe T, Gotoh K, Totoki Y, Toyoda A, Ikawa M, et al. DNA methylation of retrotransposon genes is regulated by Piwi family members MILI and MIWI2 in murine fetal testes. Genes & development. 2008;22(7):908–17. Epub 2008/04/03. doi: 22/7/908 [pii] 10.1101/gad.1640708. PubMed PMID: 18381894; PubMed Central PMCID: PMC2279202.

42. Shoji M, Tanaka T, Hosokawa M, Reuter M, Stark A, Kato Y, et al. The TDRD9-MIWI2 complex is essential for piRNA-mediated retrotransposon silencing in the mouse male germline. Developmental cell. 2009;17(6):775–87. Epub 2010/01/12. doi: S1534-5807(09)00434-1 [pii] 10.1016/j.devcel.2009.10.012. PubMed PMID: 20059948.

43. Zamudio N, Barau J, Teissandier A, Walter M, Borsos M, Servant N, et al. DNA methylation restrains transposons from adopting a chromatin signature permissive for meiotic recombination. Genes & development. 2015;29(12):1256–70. Epub 2015/06/26. doi: 10.1101/gad.257840.114. PubMed PMID: 26109049; PubMed Central PMCID: PMCPMC4495397.

44. Pastor WA, Stroud H, Nee K, Liu W, Pezic D, Manakov S, et al. MORC1 represses transposable elements in the mouse male germline. Nature communications. 2014;5:5795. Epub 2014/12/17. doi: 10.1038/ncomms6795. PubMed PMID: 25503965; PubMed Central PMCID: PMCPMC4268658.

45. Bedrosian TA, Quayle C, Novaresi N, Gage FH. Early life experience drives structural variation of neural genomes in mice. Science. 2018;359(6382):1395–9. doi: 10.1126/science.aah3378. PubMed PMID: 29567711.

46. Erwin JA, Paquola AC, Singer T, Gallina I, Novotny M, Quayle C, et al. L1-associated genomic regions are deleted in somatic cells of the healthy human brain. Nature neuroscience. 2016;19(12):1583–91. doi: 10.1038/nn.4388. PubMed PMID: 27618310; PubMed Central PMCID: PMCPMC5127747.

47. Marchetto MC, Narvaiza I, Denli AM, Benner C, Lazzarini TA, Nathanson JL, et al. Differential L1 regulation in pluripotent stem cells of humans and apes. Nature. 2013;503(7477):525–9. doi: 10.1038/nature12686. PubMed PMID: 24153179.

48. Muotri AR, Chu VT, Marchetto MC, Deng W, Moran JV, Gage FH. Somatic mosaicism in neuronal precursor cells mediated by L1 retrotransposition. Nature. 2005;435(7044):903–10. doi: 10.1038/nature03663. PubMed PMID: 15959507.

49. Richardson SR, Gerdes P, Gerhardt DJ, Sanchez-Luque FJ, Bodea GO, Munoz-Lopez M, et al. Heritable L1 retrotransposition in the mouse primordial germline and early embryo. Genome research. 2017;27(8):1395–405. doi: 10.1101/gr.219022.116. PubMed PMID: 28483779; PubMed Central PMCID: PMCPMC5538555.

50. Baillie JK, Barnett MW, Upton KR, Gerhardt DJ, Richmond TA, De Sapio F, et al. Somatic retrotransposition alters the genetic landscape of the human brain. Nature. 2011;479(7374):534–7. doi: 10.1038/nature10531. PubMed PMID: 22037309; PubMed Central PMCID: PMC3224101.

51. Evrony GD, Lee E, Park PJ, Walsh CA. Resolving rates of mutation in the brain using single-neuron genomics. eLife. 2016;5. doi: 10.7554/eLife.12966. PubMed PMID: 26901440; PubMed Central PMCID: PMCPMC4805530.

52. Evrony GD, Lee E, Mehta BK, Benjamini Y, Johnson RM, Cai X, et al. Cell lineage analysis in human brain using endogenous retroelements. Neuron. 2015;85(1):49–59. doi: 10.1016/j.neuron.2014.12.028. PubMed PMID: 25569347; PubMed Central PMCID: PMCPMC4299461.

53. Poduri A, Evrony GD, Cai X, Walsh CA. Somatic mutation, genomic variation, and neurological disease. Science. 2013;341(6141):1237758. doi: 10.1126/science.1237758. PubMed PMID: 23828942; PubMed Central PMCID: PMC3909954.

54. De Cecco M, Criscione SW, Peterson AL, Neretti N, Sedivy JM, Kreiling JA. Transposable elements become active and mobile in the genomes of aging mammalian somatic tissues. Aging (Albany NY). 2013;5(12):867–83. doi: 10.18632/aging.100621. PubMed PMID: 24323947; PubMed Central PMCID: PMCPMC3883704.

55. De Cecco M, Criscione SW, Peckham EJ, Hillenmeyer S, Hamm EA, Manivannan J, et al. Genomes of replicatively senescent cells undergo global epigenetic changes leading to gene silencing and activation of transposable elements. Aging Cell. 2013;12(2):247–56. doi: 10.1111/acel.12047. PubMed PMID: 23360310; PubMed Central PMCID: PMCPMC3618682.

56. De Cecco M, Ito T, Petrashen AP, Elias AE, Skvir NJ, Criscione SW, et al. L1 drives IFN in senescent cells and promotes age-associated inflammation. Nature. 2019;566(7742):73–8. Epub 2019/02/08. doi: 10.1038/s41586-018-0784-9. PubMed PMID: 30728521; PubMed Central PMCID: PMCPMC6519963.

57. deHaro D, Kines KJ, Sokolowski M, Dauchy RT, Streva VA, Hill SM, et al. Regulation of L1 expression and retrotransposition by melatonin and its receptor: implications for cancer risk associated with light exposure at night. Nucleic acids research. 2014;42(12):7694–707. Epub 2014/06/11. doi: 10.1093/nar/gku503. PubMed PMID: 24914052; PubMed Central PMCID: PMCPMC4081101.

58. Rodic N, Sharma R, Sharma R, Zampella J, Dai L, Taylor MS, et al. Long interspersed element-1 protein expression is a hallmark of many human cancers. Am J Pathol. 2014;184(5):1280–6. Epub 2014/03/13. doi: 10.1016/j.ajpath.2014.01.007. PubMed PMID: 24607009; PubMed Central PMCID: PMCPMC4005969.

59. Huang CR, Schneider AM, Lu Y, Niranjan T, Shen P, Robinson MA, et al. Mobile interspersed repeats are major structural variants in the human genome. Cell. 2010;141(7):1171–82. Epub 2010/07/07. doi: 10.1016/j.cell.2010.05.026. PubMed PMID: 20602999; PubMed Central PMCID: PMCPMC2943426.

60. Treiber CD, Waddell S. Resolving the prevalence of somatic transposition in Drosophila. eLife. 2017;6. doi: 10.7554/eLife.28297. PubMed PMID: 28742021; PubMed Central PMCID: PMCPMC5553932.

61. Zhuang J, Wang J, Theurkauf W, Weng Z. TEMP: a computational method for analyzing transposable element polymorphism in populations. Nucleic acids research. 2014;42(11):6826–38. doi: 10.1093/nar/gku323. PubMed PMID: 24753423; PubMed Central PMCID: PMC4066757.

62. Perrat PN, DasGupta S, Wang J, Theurkauf W, Weng Z, Rosbash M, et al. Transposition-driven genomic heterogeneity in the Drosophila brain. Science. 2013;340(6128):91–5. doi: 10.1126/science.1231965. PubMed PMID: 23559253; PubMed Central PMCID: PMC3887341.

63. Khurana JS, Wang J, Xu J, Koppetsch BS, Thomson TC, Nowosielska A, et al. Adaptation to P element transposon invasion in Drosophila melanogaster. Cell. 2011;147(7):1551–63. Epub 2011/12/27. doi: S0092-8674(11)01436-X [pii] 10.1016/j.cell.2011.11.042. PubMed PMID: 22196730; PubMed Central PMCID: PMC3246748.

64. Sytnikova YA, Rahman R, Chirn GW, Clark JP, Lau NC. Transposable element dynamics and PIWI regulation impacts lncRNA and gene expression diversity in Drosophila ovarian cell cultures. Genome research. 2014;24(12):1977–90. doi: 10.1101/gr.178129.114. PubMed PMID: 25267525; PubMed Central PMCID: PMC4248314.

65. Rahman R, Chirn GW, Kanodia A, Sytnikova YA, Brembs B, Bergman CM, et al. Unique transposon landscapes are pervasive across Drosophila melanogaster genomes. Nucleic acids research. 2015;43(22):10655–72. doi: 10.1093/nar/gkv1193. PubMed PMID: 26578579; PubMed Central PMCID: PMCPMC4678822.

66. Penke TJ, McKay DJ, Strahl BD, Matera AG, Duronio RJ. Direct interrogation of the role of H3K9 in metazoan heterochromatin function. Genes & development. 2016;30(16):1866–80. Epub 2016/08/28. doi: 10.1101/gad.286278.116. PubMed PMID: 27566777; PubMed Central PMCID: PMCPMC5024684.

67. Ninova M, Chen YA, Godneeva B, Rogers AK, Luo Y, Fejes Toth K, et al. Su(var)2-10 and the SUMO Pathway Link piRNA-Guided Target Recognition to Chromatin Silencing. Molecular cell. 2020;77(3):556–70 e6. Epub 2020/01/07. doi: 10.1016/j.molcel.2019.11.012. PubMed PMID: 31901446; PubMed Central PMCID: PMCPMC7007863.

68. Miller DE, Smith CB, Kazemi NY, Cockrell AJ, Arvanitakas AV, Blumenstiel JP, et al. Whole-Genome Analysis of Individual Meiotic Events in Drosophila melanogaster Reveals That Noncrossover Gene Conversions Are Insensitive to Interference and the Centromere Effect. Genetics. 2016;203(1):159–71. doi: 10.1534/genetics.115.186486. PubMed PMID: 26944917; PubMed Central PMCID: PMCPMC4858771.

69. mod EC, Roy S, Ernst J, Kharchenko PV, Kheradpour P, Negre N, et al. Identification of functional elements and regulatory circuits by Drosophila modENCODE. Science. 2010;330(6012):1787–97. doi: 10.1126/science.1198374. PubMed PMID: 21177974; PubMed Central PMCID: PMC3192495.

70. Rubin GM, Hong L, Brokstein P, Evans-Holm M, Frise E, Stapleton M, et al. A Drosophila complementary DNA resource. Science. 2000;287(5461):2222–4. Epub 2000/03/24. doi: 10.1126/science.287.5461.2222. PubMed PMID: 10731138.

71. Li W, Jin Y, Prazak L, Hammell M, Dubnau J. Transposable elements in TDP-43-mediated neurodegenerative disorders. PloS one. 2012;7(9):e44099. doi: 10.1371/journal.pone.0044099. PubMed PMID: 22957047; PubMed Central PMCID: PMCPMC3434193.

72. Brown EJ, Nguyen AH, Bachtrog D. The Y chromosome may contribute to sex-specific ageing in Drosophila. Nat Ecol Evol. 2020;4(6):853–62. Epub 2020/04/22. doi: 10.1038/s41559-020-1179-5. PubMed PMID: 32313175; PubMed Central PMCID: PMCPMC7274899.

73. Srivastav SP, Rahman R, Ma Q, Pierre J, Bandyopadhyay S, Lau NC. Har-P, a short P-element variant, weaponizes P-transposase to severely impair Drosophila development. eLife. 2019;8. doi: 10.7554/eLife.49948. PubMed PMID: 31845649; PubMed Central PMCID: PMCPMC6917496.

74. Wang L, Dou K, Moon S, Tan FJ, Zhang ZZ. Hijacking Oogenesis Enables Massive Propagation of LINE and Retroviral Transposons. Cell. 2018;174(5):1082–94 e12. doi: 10.1016/j.cell.2018.06.040. PubMed PMID: 30057117.

75. Okamura K, Balla S, Martin R, Liu N, Lai EC. Two distinct mechanisms generate endogenous siRNAs from bidirectional transcription in Drosophila melanogaster. Nat Struct Mol Biol. 2008;15(9):998. Epub 2008/09/05. doi: nsmb0908-998c [pii] 10.1038/nsmb0908-998c. PubMed PMID: 18769470.

76. Kawamura Y, Saito K, Kin T, Ono Y, Asai K, Sunohara T, et al. Drosophila endogenous small RNAs bind to Argonaute 2 in somatic cells. Nature. 2008;453(7196):793–7. Epub 2008/05/09. doi: nature06938 [pii] 10.1038/nature06938. PubMed PMID: 18463636.

77. Czech B, Malone CD, Zhou R, Stark A, Schlingeheyde C, Dus M, et al. An endogenous small interfering RNA pathway in Drosophila. Nature. 2008;453(7196):798–802. PubMed PMID: 18463631.

78. Brennecke J, Aravin AA, Stark A, Dus M, Kellis M, Sachidanandam R, et al. Discrete small RNA-generating loci as master regulators of transposon activity in Drosophila. Cell. 2007;128(6):1089–103. Epub 2007/03/10. doi: S0092-8674(07)00257-7 [pii] 10.1016/j.cell.2007.01.043. PubMed PMID: 17346786.

79. Saito K, Nishida KM, Mori T, Kawamura Y, Miyoshi K, Nagami T, et al. Specific association of Piwi with rasiRNAs derived from retrotransposon and heterochromatic regions in the Drosophila genome. Genes & development. 2006;20(16):2214–22. Epub 2006/08/03. doi: gad.1454806 [pii] 10.1101/gad.1454806. PubMed PMID: 16882972; PubMed Central PMCID: PMC1553205.

80. Vagin VV, Sigova A, Li C, Seitz H, Gvozdev V, Zamore PD. A distinct small RNA pathway silences selfish genetic elements in the germline. Science. 2006;313(5785):320–4. PubMed PMID: 16809489.

81. Moller HD, Mohiyuddin M, Prada-Luengo I, Sailani MR, Halling JF, Plomgaard P, et al. Circular DNA elements of chromosomal origin are common in healthy human somatic tissue. Nature communications. 2018;9(1):1069. doi: 10.1038/s41467-018-03369-8. PubMed PMID: 29540679; PubMed Central PMCID: PMCPMC5852086.

82. Shmookler Reis RJ, Lumpkin CK, Jr., McGill JR, Riabowol KT, Goldstein S. Genome alteration during in vitro and in vivo aging: amplification of extrachromosomal circular DNA molecules containing a chromosomal sequence of variable repeat frequency. Cold Spring Harbor symposia on quantitative biology. 1983;47 Pt 2:1135–9. PubMed PMID: 6574863.

83. Shibata Y, Kumar P, Layer R, Willcox S, Gagan JR, Griffith JD, et al. Extrachromosomal microDNAs and chromosomal microdeletions in normal tissues. Science. 2012;336(6077):82–6. doi: 10.1126/science.1213307. PubMed PMID: 22403181; PubMed Central PMCID: PMCPMC3703515.

84. Misra R, Shih A, Rush M, Wong E, Schmid CW. Cloned extrachromosomal circular DNA copies of the human transposable element THE-1 are related predominantly to a single type of family member. J Mol Biol. 1987;196(2):233–43. PubMed PMID: 2821286.

85. Turner KM, Deshpande V, Beyter D, Koga T, Rusert J, Lee C, et al. Extrachromosomal oncogene amplification drives tumour evolution and genetic heterogeneity. Nature. 2017;543(7643):122–5. doi: 10.1038/nature21356. PubMed PMID: 28178237; PubMed Central PMCID: PMCPMC5334176.

86. Dillon LW, Kumar P, Shibata Y, Wang YH, Willcox S, Griffith JD, et al. Production of Extrachromosomal MicroDNAs Is Linked to Mismatch Repair Pathways and Transcriptional Activity. Cell reports. 2015;11(11):1749–59. doi: 10.1016/j.celrep.2015.05.020. PubMed PMID: 26051933; PubMed Central PMCID: PMCPMC4481157.

87. Vu W, Nuzhdin S. Genetic variation of copia suppression in Drosophila melanogaster. Heredity (Edinb). 2011;106(2):207–17. doi: 10.1038/hdy.2010.41. PubMed PMID: 20606692; PubMed Central PMCID: PMCPMC3183883.

88. Flavell AJ, Ish-Horowicz D. Extrachromosomal circular copies of the eukaryotic transposable element copia in cultured Drosophila cells. Nature. 1981;292(5824):591–5. PubMed PMID: 6265802.

89. Cohen S, Yacobi K, Segal D. Extrachromosomal circular DNA of tandemly repeated genomic sequences in Drosophila. Genome research. 2003;13(6A):1133–45. doi: 10.1101/gr.907603. PubMed PMID: 12799349; PubMed Central PMCID: PMCPMC403641.

90. Mossie KG, Young MW, Varmus HE. Extrachromosomal DNA forms of copia-like transposable elements, F elements and middle repetitive DNA sequences in Drosophila melanogaster. Variation in cultured cells and embryos. J Mol Biol. 1985;182(1):31–43. PubMed PMID: 2582138.

91. Ilyin YV, Schuppe NG, Lyubomirskaya NV, Gorelova TV, Arkhipova IR. Circular copies of mobile dispersed genetic elements in cultured Drosophila melanogaster cells. Nucleic acids research. 1984;12(19):7517–31. PubMed PMID: 6093043; PubMed Central PMCID: PMCPMC320178.

92. Lanciano S, Carpentier MC, Llauro C, Jobet E, Robakowska-Hyzorek D, Lasserre E, et al. Sequencing the extrachromosomal circular mobilome reveals retrotransposon activity in plants. PLoS genetics. 2017;13(2):e1006630. doi: 10.1371/journal.pgen.1006630. PubMed PMID: 28212378; PubMed Central PMCID: PMCPMC5338827.

93. Chinen M, Lei EP. Drosophila Argonaute2 turnover is regulated by the ubiquitin proteasome pathway. Biochem Biophys Res Commun. 2017;483(3):951–7. Epub 2017/01/15. doi: 10.1016/j.bbrc.2017.01.039. PubMed PMID: 28087276; PubMed Central PMCID: PMCPMC5279065.

94. Lewis A, Berkyurek AC, Greiner A, Sawh AN, Vashisht A, Merrett S, et al. A Family of Argonaute-Interacting Proteins Gates Nuclear RNAi. Molecular cell. 2020;78(5):862–75 e8. Epub 2020/04/30. doi: 10.1016/j.molcel.2020.04.007. PubMed PMID: 32348780.

95. Clark JP, Rahman R, Yang N, Yang LH, Lau NC. Drosophila PAF1 Modulates PIWI/piRNA Silencing Capacity. Curr Biol. 2017;27(17):2718–26 e4. doi: 10.1016/j.cub.2017.07.052. PubMed PMID: 28844648; PubMed Central PMCID: PMCPMC5611886.

96. Kowalik KM, Shimada Y, Flury V, Stadler MB, Batki J, Buhler M. The Paf1 complex represses small-RNA-mediated epigenetic gene silencing. Nature. 2015;520(7546):248–52. doi: 10.1038/nature14337. PubMed PMID: 25807481; PubMed Central PMCID: PMCPMC4398878.

97. Yamanaka S, Mehta S, Reyes-Turcu FE, Zhuang F, Fuchs RT, Rong Y, et al. RNAi triggered by specialized machinery silences developmental genes and retrotransposons. Nature. 2013;493(7433):557–60. doi: 10.1038/nature11716. PubMed PMID: 23151475; PubMed Central PMCID: PMC3554839.

98. Pfeiffer BD, Ngo TT, Hibbard KL, Murphy C, Jenett A, Truman JW, et al. Refinement of tools for targeted gene expression in Drosophila. Genetics. 2010;186(2):735–55. doi: 10.1534/genetics.110.119917. PubMed PMID: 20697123; PubMed Central PMCID: PMCPMC2942869.

99. Fast I, Rosenkranz D. Temperature-dependent small RNA expression in Drosophila melanogaster. RNA biology. 2018;15(3):308–13. Epub 2018/01/19. doi: 10.1080/15476286.2018.1429881. PubMed PMID: 29345184; PubMed Central PMCID: PMCPMC5927726.

100. Kelleher ES, Jaweria J, Akoma U, Ortega L, Tang W. QTL mapping of natural variation reveals that the developmental regulator bruno reduces tolerance to P-element transposition in the Drosophila female germline. PLoS biology. 2018;16(10):e2006040. doi: 10.1371/journal.pbio.2006040. PubMed PMID: 30376574; PubMed Central PMCID: PMCPMC6207299.

101. Moon S, Cassani M, Lin YA, Wang L, Dou K, Zhang ZZ. A Robust Transposon-Endogenizing Response from Germline Stem Cells. Developmental cell. 2018;47(5):660–71 e3. doi: 10.1016/j.devcel.2018.10.011. PubMed PMID: 30393075.

102. Genenncher B, Durdevic Z, Hanna K, Zinkl D, Mobin MB, Senturk N, et al. Mutations in Cytosine-5 tRNA Methyltransferases Impact Mobile Element Expression and Genome Stability at Specific DNA Repeats. Cell reports. 2018;22(7):1861–74. Epub 2018/02/15. doi: 10.1016/j.celrep.2018.01.061. PubMed PMID: 29444437.

103. Robinow S, White K. The locus elav of Drosophila melanogaster is expressed in neurons at all developmental stages. Developmental biology. 1988;126(2):294–303. Epub 1988/04/01. doi: 10.1016/0012-1606(88)90139-x. PubMed PMID: 3127258.

104. Amarasinghe SL, Su S, Dong X, Zappia L, Ritchie ME, Gouil Q. Opportunities and challenges in long-read sequencing data analysis. Genome biology. 2020;21(1):30. Epub 2020/02/09. doi: 10.1186/s13059-020-1935-5. PubMed PMID: 32033565; PubMed Central PMCID: PMCPMC7006217.

105. Lee YS, Nakahara K, Pham JW, Kim K, He Z, Sontheimer EJ, et al. Distinct roles for Drosophila Dicer-1 and Dicer-2 in the siRNA/miRNA silencing pathways. Cell. 2004;117(1):69–81. Epub 2004/04/07. doi: S0092867404002612 [pii]. PubMed PMID: 15066283.

106. Senti KA, Jurczak D, Sachidanandam R, Brennecke J. piRNA-guided slicing of transposon transcripts enforces their transcriptional silencing via specifying the nuclear piRNA repertoire. Genes & development. 2015;29(16):1747–62. doi: 10.1101/gad.267252.115. PubMed PMID: 26302790; PubMed Central PMCID: PMCPMC4561483.

107. Belancio VP, Roy-Engel AM, Pochampally RR, Deininger P. Somatic expression of LINE-1 elements in human tissues. Nucleic acids research. 2010;38(12):3909–22. Epub 2010/03/11. doi: 10.1093/nar/gkq132. PubMed PMID: 20215437; PubMed Central PMCID: PMCPMC2896524.

108. Mumford PW, Romero MA, Osburn SC, Roberson PA, Vann CG, Mobley CB, et al. Skeletal muscle LINE-1 retrotransposon activity is upregulated in older versus younger rats. Am J Physiol Regul Integr Comp Physiol. 2019;317(3):R397–R406. Epub 2019/06/13. doi: 10.1152/ajpregu.00110.2019. PubMed PMID: 31188650.

109. Romero MA, Mumford PW, Roberson PA, Osburn SC, Parry HA, Kavazis AN, et al. Five months of voluntary wheel running downregulates skeletal muscle LINE-1 gene expression in rats. Am J Physiol Cell Physiol. 2019;317(6):C1313–C23. Epub 2019/10/17. doi: 10.1152/ajpcell.00301.2019. PubMed PMID: 31618076.

110. Nandakumar S, Grushko O, Buttitta LA. Polyploidy in the adult Drosophila brain. eLife. 2020;9. Epub 2020/08/26. doi: 10.7554/eLife.54385. PubMed PMID: 32840209; PubMed Central PMCID: PMCPMC7447450.

111. Yarosh W, Spradling AC. Incomplete replication generates somatic DNA alterations within Drosophila polytene salivary gland cells. Genes & development. 2014;28(16):1840–55. doi: 10.1101/gad.245811.114. PubMed PMID: 25128500; PubMed Central PMCID: PMC4197960.

112. Vienne J, Spann R, Guo F, Rosbash M. Age-Related Reduction of Recovery Sleep and Arousal Threshold in Drosophila. Sleep. 2016;39(8):1613–24. Epub 2016/06/17. doi: 10.5665/sleep.6032. PubMed PMID: 27306274; PubMed Central PMCID: PMCPMC4945321.

113. Ren X, Yang Z, Mao D, Chang Z, Qiao HH, Wang X, et al. Performance of the Cas9 nickase system in Drosophila melanogaster. G3 (Bethesda). 2014;4(10):1955–62. Epub 2014/08/17. doi: 10.1534/g3.114.013821. PubMed PMID: 25128437; PubMed Central PMCID: PMCPMC4199701.

114. Lee SK, Xue Y, Shen W, Zhang Y, Joo Y, Ahmad M, et al. Topoisomerase 3beta interacts with RNAi machinery to promote heterochromatin formation and transcriptional silencing in Drosophila. Nature communications. 2018;9(1):4946. Epub 2018/11/25. doi: 10.1038/s41467-018-07101-4. PubMed PMID: 30470739; PubMed Central PMCID: PMCPMC6251927.

115. Nazer E, Dale RK, Chinen M, Radmanesh B, Lei EP. Argonaute2 and LaminB modulate gene expression by controlling chromatin topology. PLoS genetics. 2018;14(3):e1007276. Epub 2018/03/13. doi: 10.1371/journal.pgen.1007276. PubMed PMID: 29529026; PubMed Central PMCID: PMCPMC5864089.

116. Hain D, Bettencourt BR, Okamura K, Csorba T, Meyer W, Jin Z, et al. Natural variation of the amino-terminal glutamine-rich domain in Drosophila argonaute2 is not associated with developmental defects. PloS one. 2010;5(12):e15264. doi: 10.1371/journal.pone.0015264. PubMed PMID: 21253006; PubMed Central PMCID: PMC3002974.

117. Zamparini AL, Davis MY, Malone CD, Vieira E, Zavadil J, Sachidanandam R, et al. Vreteno, a gonad-specific protein, is essential for germline development and primary piRNA biogenesis in Drosophila. Development. 2011;138(18):4039–50. Epub 2011/08/13. doi: dev.069187 [pii] 10.1242/dev.069187. PubMed PMID: 21831924; PubMed Central PMCID: PMC3160098.

118. Shang Y, Griffith LC, Rosbash M. Light-arousal and circadian photoreception circuits intersect at the large PDF cells of the Drosophila brain. Proceedings of the National Academy of Sciences of the United States of America. 2008;105(50):19587–94. doi: 10.1073/pnas.0809577105. PubMed PMID: 19060186; PubMed Central PMCID: PMC2596742.

119. Nagoshi E, Sugino K, Kula E, Okazaki E, Tachibana T, Nelson S, et al. Dissecting differential gene expression within the circadian neuronal circuit of Drosophila. Nature neuroscience. 2010;13(1):60–8. Epub 2009/12/08. doi: 10.1038/nn.2451. PubMed PMID: 19966839; PubMed Central PMCID: PMCPMC3878269.

120. Chirn GW, Rahman R, Sytnikova YA, Matts JA, Zeng M, Gerlach D, et al. Conserved piRNA Expression from a Distinct Set of piRNA Cluster Loci in Eutherian Mammals. PLoS genetics. 2015;11(11):e1005652. doi: 10.1371/journal.pgen.1005652. PubMed PMID: 26588211; PubMed Central PMCID: PMCPMC4654475.

121. Li H, Durbin R. Fast and accurate long-read alignment with Burrows-Wheeler transform. Bioinformatics. 2010;26(5):589–95. doi: 10.1093/bioinformatics/btp698. PubMed PMID: 20080505; PubMed Central PMCID: PMC2828108.

122. Langmead B, Trapnell C, Pop M, Salzberg SL. Ultrafast and memory-efficient alignment of short DNA sequences to the human genome. Genome biology. 2009;10(3):R25. doi: 10.1186/gb-2009-10-3-r25. PubMed PMID: 19261174; PubMed Central PMCID: PMC2690996.

